# Consequence assessment and behavioral patterns of inhibition in decision-making: modelling its underlying mechanisms

**DOI:** 10.1101/2023.02.14.528595

**Authors:** Gloria Cecchini, Michael DePass, Emre Baspinar, Marta Andujar, Surabhi Ramawat, Pierpaolo Pani, Stefano Ferraina, Alain Destexhe, Rubén Moreno-Bote, Ignasi Cos

**Affiliations:** Facultat de Matemàtiques i Informàtica, Universitat de Barcelona, Barcelona, Catalonia, Spain; Center for Brain and Cognition, DTIC, Universitat Pompeu Fabra, Barcelona, Catalonia Spain; CNRS, Paris-Saclay University, Institute of Neuroscience (NeuroPSI), Saclay, France; Department of Physiology and Pharmacology, Sapienza University of Rome, Rome, Italy; Serra-Hunter Fellow Programme, Barcelona, Catalonia, Spain

## Abstract

Learning to make adaptive decisions depends on exploring options, experiencing their consequence, and reassessing one’s strategy for the future. Although several studies have analyzed various aspects of value-based decision-making, most of them have focused on decisions in which gratification is cued and immediate. By contrast, how the brain gauges delayed consequence for decision-making remains poorly understood.

To investigate this, we designed a decision-making task in which each decision altered future options. The task was organized in groups of consecutively dependent trials, and the participants were instructed to maximize the cumulative reward value within each group. In the absence of any explicit performance feedback, the participants had to test and internally assess specific criteria to make decisions. This task was designed to specifically study how the assessment of consequence forms and influences decisions as learning progresses. We analyzed behavior results to characterize individual differences in reaction times, decision strategies, and learning rates.

We formalized this operation mathematically by means of a multi-layered decision-making model. By using a mean-field approximation, the first layer of the model described the dynamics of two populations of neurons which characterized the binary decision-making process. The other two layers modulated the decision-making policy by dynamically adapting an oversight learning mechanism. The model was validated by fitting each individual participants’ behavior and it faithfully predicted non-trivial patterns of decision-making, regardless of performance level.

These findings provided an explanation to how delayed consequence may be computed and incorporated into the neural dynamics of decision-making, and to how learning occurs in the absence of explicit feedback.

## 1 INTRODUCTION

The brain mechanisms involved in taking decisions are of key interest and have been studied extensively in the last decades (reviewed in (1,2)). Many studies focused on characterizing the neural dynamics of reward processing (3–5), visual discrimination (6–8), and multiple other aspects involved in option assessment during value-based decision-making (9–12). Other tasks were developed to study decisions in the context of short-term memory (13), and cost-risk trade-off (14–16). In most of these contexts, choice outcomes are immediately feedbacked. This feature makes calculating costs and benefits straightforward, as all the necessary information is directly available to the decision maker for calculation (17–20). However, a complete account of value-based choice behavior requires understanding how the brain detects and computes the non-immediate consequences of choices, and how to use this information to guide subsequent decision strategies. Despite the rich literature in cognitive decision-making and the fact that long-term consequence is a critical concern in our daily decision-making processes, the dynamics of its operation have not yet been incorporated into state-of-the-art models of decision-making (21–23). These classic models, by construction, work for independent consecutive trials by considering accumulation of evidence about choice alternatives for each trial separately (24), but they do not take into consideration neither the memory of recent past nor the long-term effects of decisions. By contrast, studies on hierarchical decision-making investigate the case when the choice made in one trial influences the available choice alternatives in the sub-sequent one (25). In this scenario, the case when the immediate most rewarding choice leads to a lower long-term reward is of particular interest. Moreover, if this relationship is latent, what are the cognitive mechanisms that make us learn the optimal strategy? Furthermore, how does the learning occur in the absence of an explicit performance feedback?

To answer these questions, in this manuscript we studied a specific case of decision-making that we called consequence based. Namely, we organized consecutive perceptual decision-making trials into groups of dependent trials, where the choice made in one trial has a consequence on the next one by determining the available choice options. How does the complexity of a perceptual decision-making task augment when combined with consequence assessment? First, consequence-based decisions require an increased temporal span of consideration, and, consequently, involve a more uncertain and broader set of factors to examine. This typically results in more computationally demanding option evaluation (26–29), longer deliberation, and often poorer decision accuracy (30,31). Second, making decisions based on gauging choice consequence involves a range of cognitive processes broader than those involved in immediate sensory-motor decisions (32,33), with particular emphasis on value integration (34,35), metacognitive processing (36,37) and long-term working memory (38,39). Despite the fact long-term consequence assessment may be viewed as a time extended version of immediate actions’ outcomes evaluation, significant doubts remain regarding the core cognitive and neural processes underlying this capability (40).

To investigate the cognitive processes underlying consequence-based decision-making, we carried out a combined experimental and theoretical study. In the first part of this work, we designed a behavioral paradigm, the *consequential task*, to characterize consequence-based option assessment in a decision-making context in which there was no explicit performance feedback. In this task, trials were organized in groups (of one, two, or three trials). At each trial, participants were instructed to make a binary choice between two stimuli (each associated with a specific reward quantity) with the goal to find the strategy that yielded the most cumulative reward value across each group of trials. In brief, the selected stimulus in a trial influenced the available stimuli presented in the next trial in such a way that choosing a large/small stimulus would yield lower/higher average stimuli in the next trial. Crucially, these changes were neither cued nor part of the instruction given to the participants at the beginning of the session. Moreover, no performance metric was provided to the participants, who, therefore, had to rely on their own internal assessment of performance based on the implicit changes caused by their choices on the stimuli of subsequent trials. To summarize, the consequential task implemented consequence through linked trials, its nature was not disclosed in the instructions given to the participants, and no explicit performance feedback was provided.

The consequential task combined features common both to hierarchical decision-making (41–43) and delay discounting paradigms (44–48). However, the absence of an explicit performance metric during the task made our paradigm particularly suitable to study how learning the optimal strategy may evolve from shortsighted to long-term predictions of the next states. In contrast to standard hierarchical decision-making and partially observable Markov decision processes (49,50), in the consequential task participants were not aware of the underlying structure. Participants were informed that the choice they made in one trial had an influence on the following one, but it was not clearly stated that their action led to mutually exclusive states, i.e., the possible scenarios a participant can encounter in a group of trials. Moreover, participants did not know if they had found the optimal solution, i.e., picked the correct sequence of decisions to maximize cumulative reward value. On the other hand, delay discounting tasks focused on the principle of inhibitory short-term control where the presence of explicit cues helped overcoming impulsive behavior, such as in the marshmallow experiment (51,52), or the *farming on Mars* task (53) (see review in the Discussion). By contrast, the purpose of our study was to understand how participants learned the effects of their actions on the environment, as opposed to assessing whether reward value varies with time. In other words, the absence of explicit learning cues was intended to force the participants to rely on their own subjective performance feedback to infer the delayed consequence of their decisions across groups of successive trials.

In the second part of our study, we provided a theoretical framework of the cognitive and neural processes required for consequence-based decision-making, including the patterns of inhibition and of far-sighted consequence assessment instrumental to gain the most reward across trials. The model was organized in three layers. The bottom layer, in line with the Amari, Wilson-Cowan and Wong-Wang models (21,54–58), described the neural dynamics of binary decision-making by means of two populations of neurons. The middle and top layers implemented an oversight mechanism for the assessment of consequence across groups of trials and the learning mechanism as a function of an objective perception of reward value across trials. This model reproduced the full variety of behavioral observations across the different participants accurately while predicting a plausible neural implementation of the processes underlying the learning of consequence-based decision-making. In particular, our model described how the metacognitive assessment of consequence extends from shortsighted to long-term value prediction through an oversight mechanism that monitors predicted performance.

## 2 RESULTS

### 2.1 Task design

In this section, we describe the consequential task, specifically designed to tap into the cognitive mechanisms involved in learning delayed consequences in the absence of performance feedback. In this task, 28 healthy participants were instructed to choose one of the two stimuli, depicting reward values through differently filled water containers, presented left and right on the screen. The participants reported their choices by sliding the computer mouse’s cursor from the central cue to the chosen stimulus (see Figure 1 and Materials and Methods for a thorough description).

**Figure 1.**
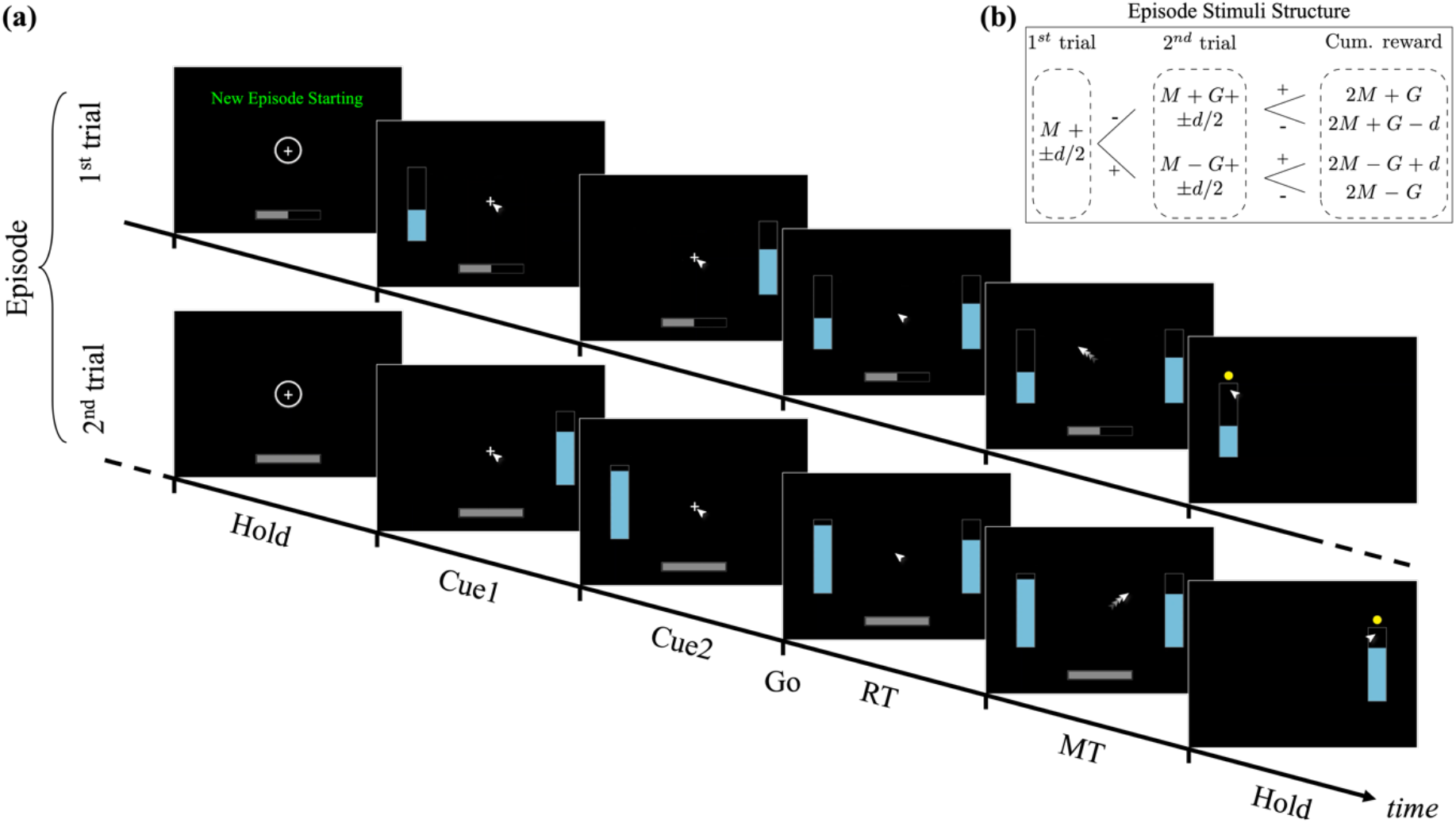
Time-course of a typical horizon 1 episode of the consequential decision-making task. **(a)** The episode consists of two dependent trials. The first starts with the message “New Episode Starting” in the center-top of the screen, a circle surrounding a cross in the center (central target), and half full progress bar at the bottom of the screen. The progress bar indicates the current trial within the episode (for horizon 1, 50% during the first trial, 100% during the second trial). After holding for 500ms, the left or right (chosen at random) stimulus is shown, followed by its complementary stimulus 500ms later. Both stimuli are shown together 500ms later which serves as the GO signal. At GO, the participant has to slide the mouse from the central target to the bar of their choosing. Once the selected target is reached, a yellow dot appears over that target. The second trial follows the same pattern as the first. See Methods for more details. **(b)**: Construction scheme for the size of the stimuli in each episode. The first trial within the episode consists of 2 stimuli of size M+d/2 and M-d/2. The second trial within the episode depends on the selection made in the previous trial. If the first selected stimulus is M-d/2 (following symbol “-” in the figure), then the second trial consists of stimuli with size M+G+d/2 and M+G-d/2, otherwise M-G+d/2 and M-G-d/2 (following symbol “+” in the figure). The cumulative reward value of the episode can therefore assume 4 distinct values (ordered from best to worst): 2M+G, 2M+G-d, 2M-G+d, and 2M-G. See Methods for more details on the values of M, G, d.

Since consequence depends on a predictive assessment of future contexts, the task was organized into two types of trial blocks, in which the participants had to maximize the reward value. There were blocks in which trials required one-shot decisions, purely independent from each other. As in most typical decision-making paradigms, the reward value in these trials could be maximized by picking the best available option in that instance. However, in other blocks, trials were grouped into pairs or triads of dependent trials. We called each group of consecutive trials an episode to signify the boundary of dependence between them, and defined the notion of horizon (*n_H_*) as a metric for the depth of consequence to be expected for that episode. The horizon of a specific episode equaled the number of dependent trials following the first trial of each episode. For example, for *n_H_*=1 an episode consists of 2 trials. The nature of the dependence between trials of an episode was such that the mean reward values of the stimuli in the second/third trial were systematically increased or decreased based on the participant’s choice in the preceding trial. Specifically, choosing the larger stimulus value led to a reduction of stimuli values in the subsequent trial, whereas achieving greater future value options required deliberately choosing the lesser option in the previous trial (Figure 1b). The increment/reduction amount (*G*) was set constant and chosen such that selecting the larger stimulus could never compensate for the loss. In other words, the optimal performance across the task was achieved by choosing “big” in single trial episodes (horizon *n_H_*=0), and deliberately choosing “small” in all trials of *n_H_*=1 and *n_H_*=2 episodes except the last, in which “big” should be chosen.

Participants were instructed that their goal was to maximize the cumulative reward value per episode. Learning the optimal policy was made challenging by a number of different factors. First, perceptual discrimination, quantifying the size difference between stimuli varies within 1-20% of the container. Second, although the participants were instructed that their choices may affect future trials within the episode, the nature of this dependency was not signaled in any obvious way. This means that from the perspective of the participants, the value of the reward offers might at first appear random. Third, explicit performance feedback after each episode was crucially omitted from the task. The reason for this is that the presence of performance feedback might have had the undesirable effect of participants focusing on finding the specific sequence of choices within episode yielding optimal performance feedback, without having to learn the relationship between their decisions and the subsequent trials. In other words, an explicit measure of performance might have reduced the task to an explicit trial-and-error test of deciding for example, “big-small”, “small-big”, etc., until finding the sequence of choices leading to maximum performance, rather than learning to evaluate each option’s consequence in terms of their prediction of future reward value to attain the goal. In contrast, the absence of performance feedback made the participants not informed about their performance throughout the block, and ought to oblige them to create an internal sense of assessment, which can only rely on two mechanisms: the sensory perception of the systematic stimuli changes in the subsequent trial after each choice, and the exploration of option choices at each trial during the earlier part of each block. The resulting task essentially becomes a measure of learning about delayed consequences associated with each option in the absence of explicit performance feedback.

In summary, for the participants to be able to perform the task, they were informed of the episode-based organization of trials at each block, i.e., the horizon. The instruction to the participant was to find the strategy leading to the most cumulative reward value for each episode and, for the reasons mentioned previously, to actively explore their choices. Further details are shown in the Methods section, and in Figure 1.

### 2.2 Behavioral Results

The metrics extracted from the participants’ behavioral data were their performance (PF), reported choices (CH), reaction time (RT), and visual discrimination (VD) sensitivity. The PF is a single-episode metric assuming values from 0 (worst) to 1 (best); it is calculated as the percentage of reward value obtained throughout the episode normalized by the maximum and minimum that could have been obtained. CH was the choice made by the participant in each trial, in terms of small or large reward stimulus. The RT was calculated as the time difference between the simultaneous presentation of both stimuli (the GO signal), and the onset of the movement. The VD is the ability to visually discriminate between stimuli, i.e., identifying which one is the bigger/smaller (see Methods for further details). As shown below, when the difference between stimuli (DbS) is small, participants were not able to accurately distinguish between stimuli. The DbS varies within 1-20% of the size of the container.

The absence of explicit performance-related feedback at the end of each episode made the task more difficult, and, consequently, not all participants were able to find the optimal strategy. For horizon *n_H_=0,* all 28 participants but two learned and applied the optimal strategy, i.e., repeatedly selecting the larger stimulus. By contrast, only 22 participants learned the optimal strategy during horizon *n_H_=1,2* blocks, i.e., selecting the larger stimulus in the last trial only.

We analyzed the exploratory strategy participants used. In particular, we tested whether participants only considered the size of the stimuli (small/big), or if they also tried other hypotheses, such as the order of presentation (first/second) or the location (left/right) of the stimuli. The result of this analysis can be found in the Supplementary Materials (FigSupp 1). In brief, participants mostly considered only the size as a possible factor for optimization. Most participants who did not learn the optimal strategy for *n_H_=1,2* repeatedly chose the larger stimulus for all trials.

In Materials and Methods, Sec. Consequential Decision-Making task, we described how the task was structured, and we mentioned that we randomized the order in which participants performed the horizons. This means that, for example, some participants performed *n_H_* 2 before *n_H_* 0. We wondered if the order of the execution of the horizons had an influence on the learning. To address this, we performed an analysis comparing learning times on different conditions. The results of this investigation can be found in the Supplementary Materials (FigSupp 2 and FigSupp 3). In brief, we discovered that once the optimal strategy was understood in *n_H_* 1 or *n_H_* 2, participants generalized the rule and by abstraction applied it to the horizon performed afterwards. For this reason, we defined a single learning time per participant which refers to the whole session. In other words, we called *learning time (t_L_)* the first episode (*E*) of the session in which the optimal strategy was assimilated. Namely, we defined the time at which the strategy was assimilated as the moment after which the optimal strategy was used in at least 9 out of the following 10 episodes, and 75% of the remaining episodes until the end of the block. To ensure that a low success rate was not caused by perceptual discrimination errors (during low VD), we excluded the most difficult episodes in terms of DbS to calculate the learning time.

Figure 2 shows the summary results for all 28 participants. In Panel (a), we show the histogram of their learning times in terms of episodes (*E*). The last histogram bar in Figure 2a (shown as NL – No Learning), shows the aggregate of the 6 participants who never learned the optimal strategy. We can identify four types of participants as a function of their learning speed: slow, medium, fast learners, and those participants who did not ever learn the strategy.

Figure 2b shows the VD, for all difficult trials (smallest DbS) and participants, where VD was calculated as the percentage of correct choices over the last 80 episodes in the horizon *n_H_=0* block. On average, stimuli were discriminated correctly in 71% of the most difficult trials. Thus, despite having learned the optimal strategy, because of the low VD, most participants continued making some errors. This is reported in Figure 2c, showing the grand average and standard error of the PF across subjects as a function of the difficulty level of the episode, for all episodes following each participant’s learning time (*p*=10^-12^, F-stat=59). Note that, in Figure 2d, the RT gradually increased with growing difficulty to discriminate the stimuli, thus exhibiting a gradual and significant sensitivity to VD (*p*=10^-25^, F-stat=160).

**Figure 2.**
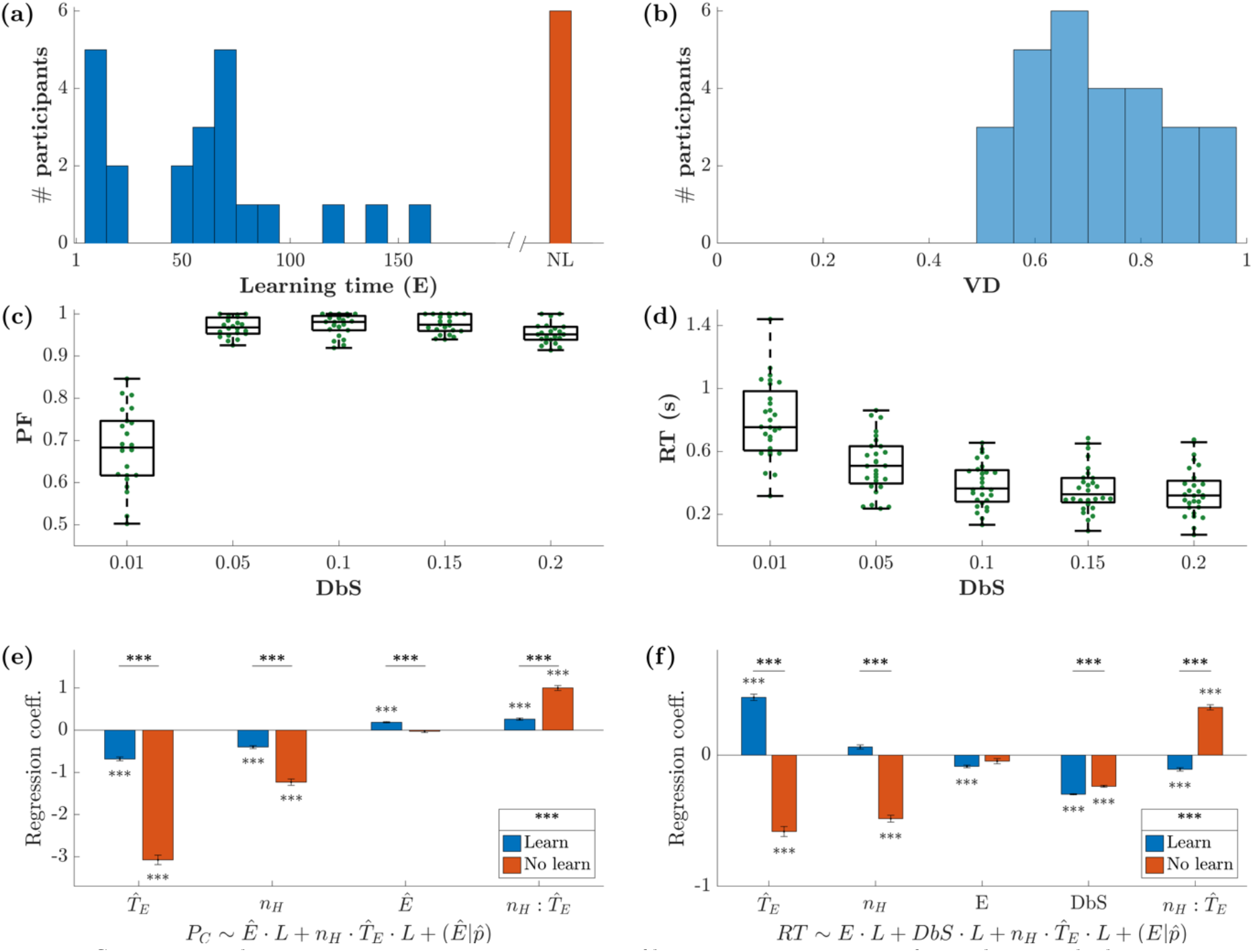
Summary results across participants. **(a)** Histogram of learning times, in terms of episodes (E). The learning time is defined as the first episode throughout the whole session in which the optimal strategy was applied repeatedly (see Methods). We identified four groups of participants: fast, medium and slow learners, and participants who did not discover the optimal strategy (NL – No Learning). **(b)** Histogram of the visual discrimination (VD) calculated by computing the percentage of correct selections of the last 80 episodes, in the horizon 0 block, for only the most difficult trials (DbS d=0.01). **(c)** Performance as a function of DbS, for the trials after the optimal strategy was applied. **(d)** Reaction Time (RT) versus DbS. The more similar the stimuli, the longer participants needed to make a decision. (e-f) Regression coefficients for the linear mixed-effects models *P*_*oc*_∼*L* ⋅ *Ê* + *L* ⋅ *n*_#_ ⋅ *T*_*E*_ + *Ê*+*p^*) and *RT*∼*L* ⋅ *Ê* + *L* ⋅ *d* + *L* ⋅ *n*_#_ ⋅ *T_E_* + (*Ê*|*p^*), where *P*_*oc*_ is the percentage of optimal choices, RT is the reaction time, E is the episode number, *Ê* counts episodes in groups of 10, *n*_#_ is the horizon number, *T*_*E*_ is the trial within episode counting backwards from last to first, d is the DbS, *n*_#_: *T*_*E*_ is the interaction term, and *p^* is the participant. We used maximum likelihood to estimate the model parameters. Participants were divided into two groups: those who learned the optimal strategy (blue) and those who did not (red), see Panel (a). The statistical difference between learning groups in reported next to the legend.

The dependency of PF and RT on VD together with the other variables must be established statistically. To assess the learning process, we quantified the relationship of PF and RT with horizon *n_H_*, trial within episode *T_E_*, and episode *E*. To obtain consistent results, we adjusted these variables as follows. The trial within episode is reversed, from last to first, because the optimal choice for the last *T_E_* (large) is the same regardless of the horizon number. The variable representing the trial within episode counted backwards is denoted as *Ť*. Furthermore, regarding the model for PF, to consider trials within episode independently, we adapted the notion of PF (defined as a summary measure per episode) to an equivalent of PF per trial, i.e., the percentage of optimal choices *P*_*oc*_. To be able to calculate such percentage, we grouped the episodes in blocks of 10 and used their average. This new variable is called *Ê*. Regarding the model for RT, since we consider each episode separately, and not an aggregate of 10 of them, we also check the dependency with DbS (*d*). Finally, to assess the difference between learning groups, we introduce the categorical variable L that identifies the group of participants that learned the optimal strategy and the ones who did not, according to Figure 2a. We then used a linear mixed effects model (59,60) to predict PF and RT. The independent variables for the fixed effects are horizon *n_H_*, trial within episode *Ť*! (counted backwards), and the passage of time expressed as groups of 10 episodes *Ê* each for PF, or for RT the episode *E* and DbS *d*. We set the random effects for the intercept and the episodes grouped by participant *p^*; we write the random effects as ’*Ê*(*p^*). The resulting models are: *P*_*oc*_∼*L* ⋅ *Ê* + *L* ⋅ *n*_$_ ⋅ *Ť* + ’*Ê* (*p^*) and *RŤ*∼*L* ⋅ *Ê* + *L* ⋅ *d* + *L* ⋅ *n*_$_ ⋅ *Ť* + (*Ê*|*p^*). The regression coefficients, with their respective group significance, are shown in Figure 2e-f. The detailed results of the statistical analysis are reported in the Supplementary Materials (Table 2). In panel (e), *P*_*oc*_decreases with *Ť*, suggesting that the first trial(s) within the episode are less likely to be guessed right, i.e., favoring the smaller of both stimuli. This makes sense, since only the early trials within the episode required inhibition. Moreover, looking at the amplitude of the regression coefficients, we can state that this has a larger impact in the no-learning case. The same argument can be made for the dependency with *n_H_*. The mayor difference between learning and no-learning can be appreciated when considering the time dependence: for the learners’ group *P*_*oc*_ increases as time goes by, i.e., *Ê* increases, while it is not significant for the group that did not learn the optimal strategy. The two learning groups are globally statistically different (*p=10^-12^*). In panel (f), RT shows converse effect directions between learning and no-learning groups for both dependencies on *Ť*! and *n_H_*. The participants who learned the optimal strategy exhibited longer RT for the earlier trials within the episode, consistently with the need of inhibiting the selection of the larger stimulus. Also, the larger the horizon, the longer the RT, opposite to the no-learning group. As expected, RT increases when decreasing DbS for both groups. The two learning groups are globally statistically different (*p=10^-17^*).

**Table 1.**
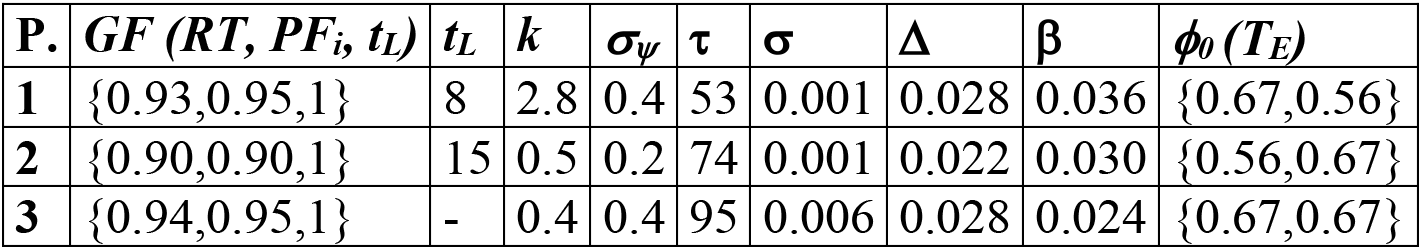
Parameter values obtained when fitting data from 1 block for each of the 3 participants. The parameters 1, τ, σ, Δ α, and β refer to Eq. 3; <0 and k belong to Eq. 4; α_ψ_ is deployed in Eq. 2. The learning time (tL) and the goodness of fit (GF) are shown in the first 2 columns.

| P. | $GF(RT, PF_i, t_L)$ | $t_L$ | $k$ | $\sigma_\psi$ | $\tau$ | $\sigma$ | $\Delta$ | $\beta$ | $\phi_0(T_E)$ |
| --- | --- | --- | --- | --- | --- | --- | --- | --- | --- |
| 1 | {0.93,0.95,1} | 8 | 2.8 | 0.4 | 53 | 0.001 | 0.028 | 0.036 | {0.67,0.56} |
| 2 | {0.90,0.90,1} | 15 | 0.5 | 0.2 | 74 | 0.001 | 0.022 | 0.030 | {0.56,0.67} |
| 3 | {0.94,0.95,1} | - | 0.4 | 0.4 | 95 | 0.006 | 0.028 | 0.024 | {0.67,0.67} |

**Table 2.** Linear mixed effects model for the percentage of optimal choices selected *P*_*oc*_ and for the reaction time *RT*. The independent variables for the fixed effects are horizon n_H_, trial within episode *Ť*_*E*_ (counted backwards), and the passage of time expressed as groups of 10 episodes *Ê* each for PF, or for RT the episode E and DbS d. We set the random effects for the intercept and the episodes grouped by participant *p^*.

| | $P_{oc} \sim L \cdot \hat{E} + L \cdot n_H \cdot \hat{T}_E + (\hat{E}\vert \hat{p})$ | | | | | | $RT \sim L \cdot E + L \cdot d + L \cdot n_H \cdot \hat{T}_E + (E\vert \hat{p})$ | | | | | |
| --- | --- | --- | --- | --- | --- | --- | --- | --- | --- | --- | --- | --- |
| F-stat. | 139.7 |  |  |  |  |  | 205.9 |  |  |  |  |  |
| p-value | 0 |  |  |  |  |  | 0 |  |  |  |  |  |
| Fixed effects | Estimate | SE | tStat | pVal | Lower | Upper | Estimate | SE | tStat | pVal | Lower | Upper |
| Intercept | 3.38 | 0.25 | 13.7 | $10^{-40}$ | 2.89 | 3.86 | 0.75 | 0.15 | 4.95 | $10^{-07}$ | 0.456 | 1.05 |
| $\hat{T}_E$ | -3.07 | 0.19 | -16.2 | $10^{-55}$ | -3.45 | -2.70 | -0.58 | 0.08 | -7.04 | $10^{-12}$ | -0.75 | -0.42 |
| $n_H$ | -1.23 | 0.13 | -9.30 | $10^{-20}$ | -1.49 | -0.97 | -0.48 | 0.06 | -8.36 | $10^{-17}$ | -0.60 | -0.37 |
| $\hat{E}$ | -0.03 | 0.04 | -0.68 | 0.5 | -0.11 | 0.05 | - | - | - | - | - | - |
| $E$ | - | - | - | - | - | - | -0.05 | 0.04 | -1.21 | 0.23 | -0.13 | 0.03 |
| $d$ | - | - | - | - | - | - | -0.24 | 0.02 | -15.78 | $10^{-55}$ | -0.27 | -0.21 |
| $L_1$ | -1.96 | 0.28 | -7.02 | $10^{-12}$ | -2.50 | -1.41 | -1.45 | 0.17 | -8.42 | $10^{-17}$ | -1.79 | -1.11 |
| $\hat{T}_E:n_H$ | 0.99 | 0.10 | 9.88 | $10^{-22}$ | 0.80 | 1.20 | 0.36 | 0.04 | 8.21 | $10^{-16}$ | 0.28 | 0.45 |
| $\hat{T}_E:L_1$ | 2.28 | 0.21 | 10.66 | $10^{-25}$ | 1.86 | 2.70 | 1.11 | 0.09 | 11.89 | $10^{-32}$ | 0.93 | 1.30 |
| $n_H:L_1$ | 0.78 | 0.15 | 5.22 | $10^{-07}$ | 0.49 | 1.07 | 0.62 | 0.07 | 9.48 | $10^{-21}$ | 0.49 | 0.75 |
| $\hat{E}:L_1$ | 0.20 | 0.047 | 4.29 | $10^{-05}$ | 0.11 | 0.30 | - | - | - | - | - | - |
| $E:L_1$ | - | - | - | - | - | - | -0.02 | 0.04 | -0.49 | 0.62 | -0.11 | 0.07 |
| $d:L_1$ | - | - | - | - | - | - | -0.06 | 0.02 | -3.42 | $10^{-3}$ | -0.09 | -0.02 |
| $\hat{T}_E:n_H:L_1$ | -0.69 | 0.11 | -6.11 | $10^{-09}$ | -0.92 | -0.47 | -0.51 | 0.05 | -10.31 | $10^{-25}$ | -0.61 | -0.42 |

Out of all 28 participants we analyzed, in Figure 3 we show the data from 3 sample participants. Figure 3 shows their associated PFs, CHs, and RTs metrics, and the order of execution of the different blocks and horizons. Each column corresponds to a participant and each row to a different horizon level. Note that all three participants performed the *n_H_=0* task correctly (Figure 3a,b). The first 2 participants also performed *n_H_=1* correctly, while participant 3 did not learn the correct strategy until he executed *n_H_=2*. Note that participants 1 and 2 performed *n_H_=1* before *n_H_=2*, they learned during *n_H_=1,* and then applied the same strategy for *n_H_=2*. Because of this, a very fast learning process can be noted during the first *n_H_=2* block. In Figure 3c, note that some RTs are negative. In these cases, the participant did not wait for the presentation of the GO signal to start the movement.

**Figure 3.**
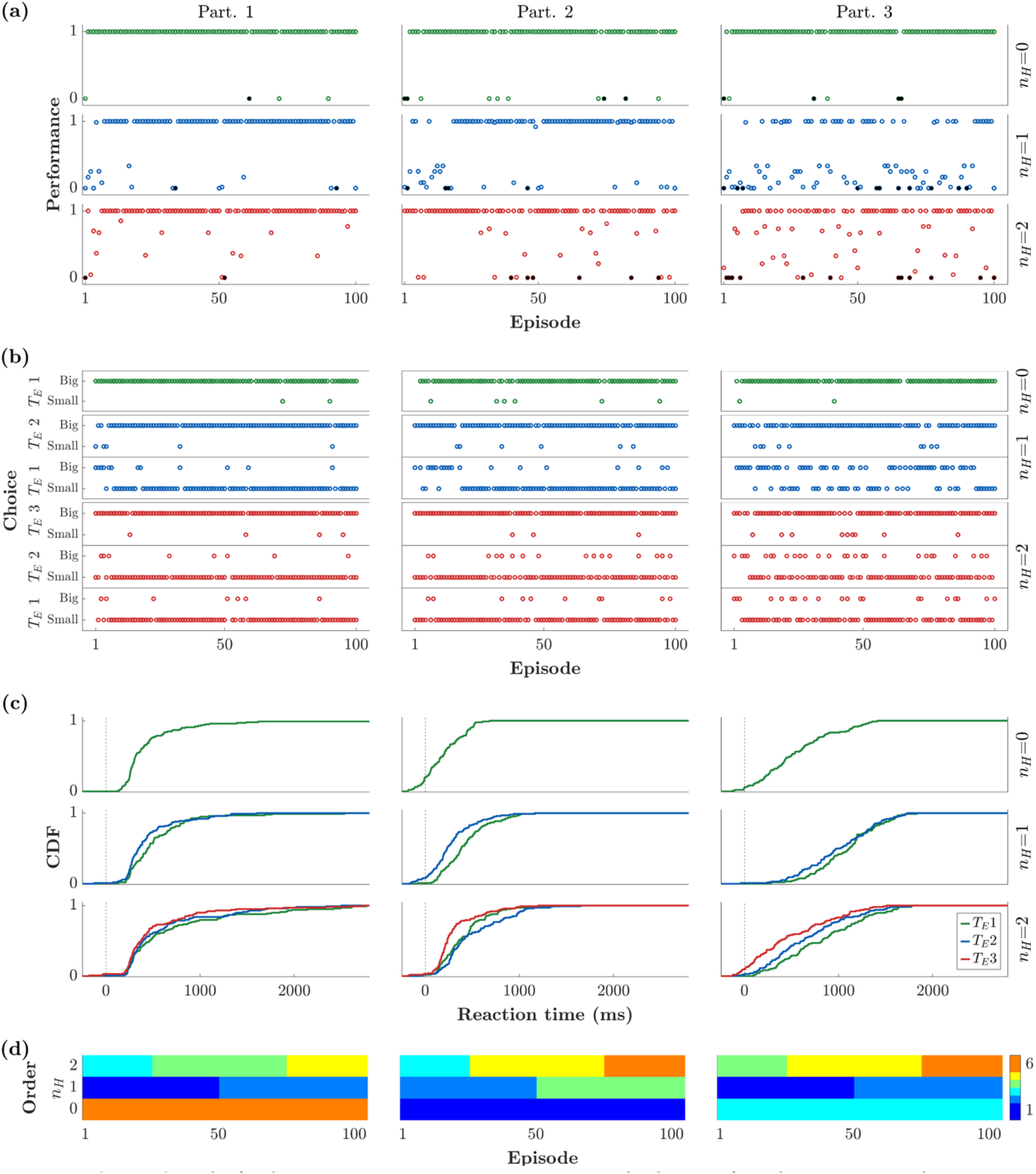
Behavioral results for three representative participants. Rows and columns refer to horizons (nH) and participants, respectively. (a) Performance per episode. (b) Choice behavior per trial, in terms of selecting the bigger or smaller stimulus. Results are gathered by horizon (nH) and respective trial within episode (TE). (c) Cumulative density function (CDF) of reaction times. The color code indicates the trial within episode (green for TE=1, blue for TE=2, and red for TE=3). (d) Order of execution of blocks and horizons.

### A Neurally-inspired Model of Consequential Decision-Making

In this section, we describe our mathematical formalization of consequential decision-making, incorporating a variable foresight mechanism, adaptive to the specifics of how reward is distributed across trials of each episode. We formalized these processes using a three-layer neural model, described next. In brief, we used a mean-field model for binary decision-making, driven by a system able to learn the optimal strategy, and consequently dictate the choices to the decision-making process. The reason why we chose to build such a model instead of employing, for example, a classic reinforcement learning model is that our model not only describes behavioral patterns of learning, but it is also biophysically plausible. The neural dynamics in the mean-field approximation have been derived analytically from a network of spiking neurons used for making binary decisions (61).

#### 2.3.1 Layer 1: Neural dynamics

To describe the neural dynamics at each trial, we used a mean-field approximation of a biophysically based binary decision-making model (23,58,61,62). This approximation has been often used to analytically study neuronal dynamics, through analysis of population averages. This included a simplified version that reproduced most features of the original spiking neuron model while using only two internal variables (21).

The core of the model consists of two populations of excitatory neurons: one sensitive to the stimulus on the left-hand side of the screen (L), and the other to the stimulus on the right (R). The intensity of the evidence is the size of each stimulus, which is directly proportional to the amount of reward displayed. In the model this is captured by the parameters λ_L_, λ_R,_ respectively. Although in the interest of our task we distinguish between the bigger and smaller stimulus values, in the formulation of the model it is convenient to characterize stimuli based on their position, i.e., left/right. The reason here is that the information on which target is bigger is already conveyed by the respective stimuli values, i.e., the parameters λ_L_, λ_R_. Moreover, this allows to introduce an extra degree of freedom in the model, without increasing the number of variables. The equations

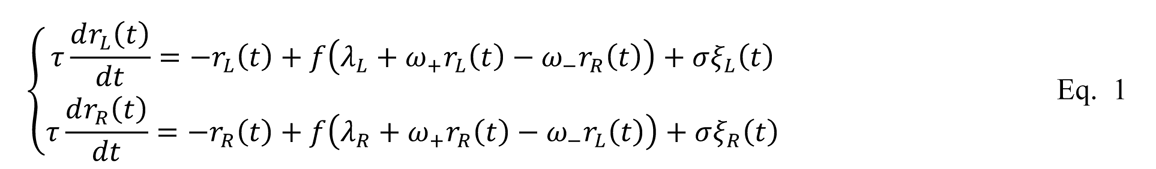

describe the temporal dynamics of the firing rates (*r_L_, r_R_*) for each of the two populations, and may be interpreted as originating from a neural network as shown in Figure 4a. Each pool has recurrent excitation (ω_+_), and mutual inhibition (ω_-_). Although the schematic indicates that both excitation and inhibition emanate from a single population of excitatory neurons, this connectivity could be achieved with an equivalent network of excitatory and inhibitory subpopulations (21,22,55,62,63). In particular, we refer to the work by Wong and Wang (21), where they reduced a spiking neural network of both excitatory and inhibitory neurons to a two-variable system describing the firing rate of the mean-field dynamics of two populations of excitatory neurons. We opted for this simplified architecture because they are equivalent under some conditions and provide a more compact formulation. Furthermore, the network shares a basic feature with many other models of bi-stability: to ensure that only one population is active at any time (mutual exclusivity; (64,65)), mutual inhibition is exerted between the two populations ((66–68)). The overall neuronal dynamics are regulated by the time constant τ, and Gaussian noise ξ with zero mean and standard deviation σ. The sigmoidal function *f* is defined as *f(x)* = *f*_)*max*_IA1 + *exp*−(*x* − *θ*)⁄*k*^F^<H, with *F*_)*+_ denoting the firing rate saturation value.

**Figure 4.**
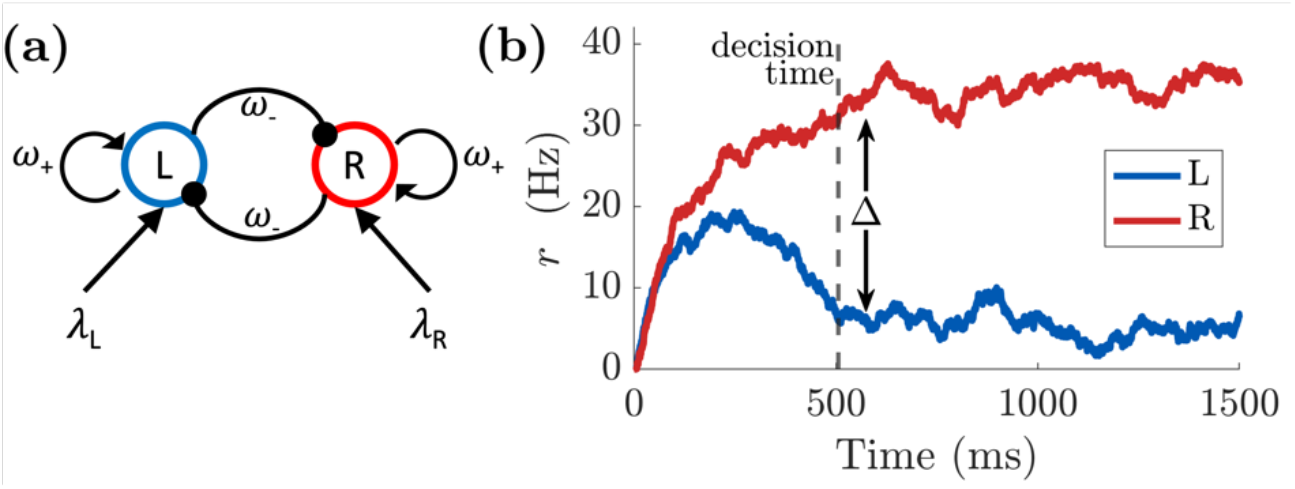
(a) Network structure of binary decision model of mean-field dynamics. The L pool is selective for the stimulus L (λL), while the other population is sensitive to the appearance of the stimulus R (λR). The two pools mutually inhibit each other (ω-) and have self-excitatory recurrent connections (ω+). **(b)** Firing rate of the two populations (L, R) of excitatory neurons according to the dynamics in *Eq. 1*. A decision is taken at time 506 ms (vertical dashed line) when the difference in activity between L and R pools passes the threshold of Δ =25Hz. The strengths of the stimuli are set to λL = 0.0203 and λR = 0.0227. The time constant and the noise are set to τ = 80 ms and σ = 0.003 ms^-1^, respectively.

The neural dynamics described in this section refer to the time-course of a single trial, and is related to the discrimination of the two stimuli. The model commits to a perceptual decision when the difference between the L and R pool activity crosses a threshold Δ (69), see Figure 4b. This event defines the trial’s decision time. Note that the decision time and the likelihood of picking the larger stimulus are conditioned by the evidence associated with the two stimuli (β_L_, β_R_), i.e., how easy it is to distinguish between them. Namely, the larger the difference between the stimuli is, the more likely and quickly it is that the larger stimulus is selected.

This type of decision-making model is made such that the larger stimulus is always favored. Although the target with the stronger evidence in Eq. 1 is the most likely to be selected, this behavior becomes a particular case when this first layer interacts with the middle layer of our model, as described in the next section.

#### 2.3.2 Layer 2: Intended decision

While most decision-making models consider only information involving one-shot decisions (21,69–72), the increased temporal span consideration and the uncertainty due to the consequence of the decision-making processes involved in the consequential task require additional elements for our model. The second layer of our model is devoted to build a mechanism capable of dynamically shifting from the natural (perceptual based) impulse of choosing the larger stimulus, to inhibiting that preference and choosing the smaller one. We implemented such a mechanism by means of an inhibitory control pool, which regulates, when desired, the reversal of the selection criterion towards the smaller or larger stimulus. We called this mechanism *intended decision*, as it defines the intended target to select at each trial. This constitutes the layer enabling the model to switch preference as a function of the context (see layer 3 description).

Specifically, the intended decision mechanism at each trial is represented as a two-attractor dynamical system. If the state of the model may be interpreted as the continuous expression of its tendency for one over another choice, an attractor is the state towards which the dynamics of the system naturally evolve. Since we have two choices, to implement this we considered the energy function *Ê*(ψ) = ψ,(ψ − 1), that has two basins of attraction at 0 and 1, associated to the small and big stimulus, respectively (see Figure 5a). Hence, the dynamics of *ψ* are regulated by

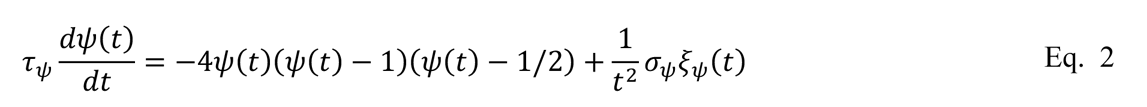

where τ*_ψ_* is a time constant. The Gaussian noise *ξ_ψ_(t)* is scaled by a constant (*σ_ψ_*) and decays quadratically with time. Thus, the noise exerts a strong influence at the beginning of the process and becomes negligible as one of both basins of attraction is reached.

**Figure 5.**
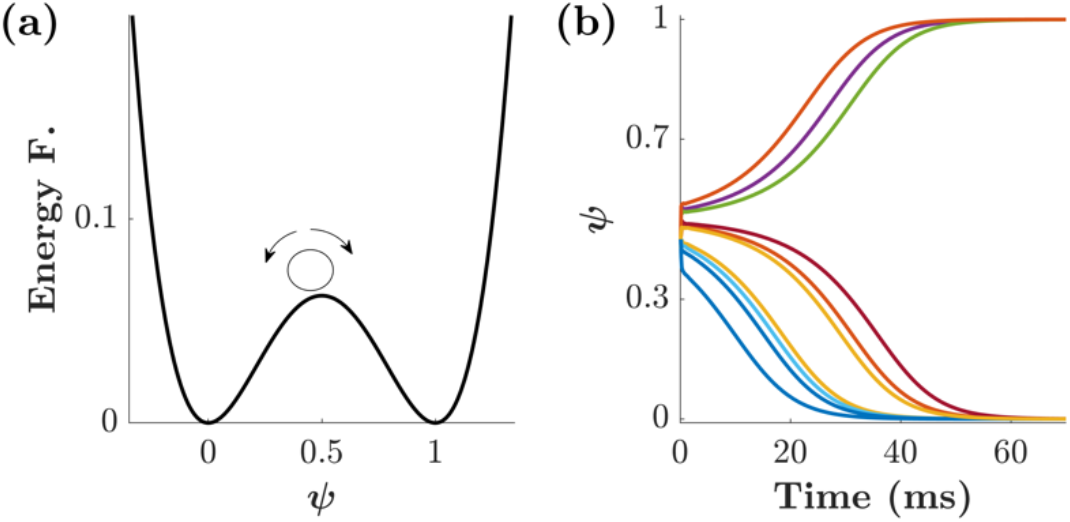
Dynamics of the second layer of the model. a) Energy function *E*(*ψ*) = *ψ*^2^(*ψ* − 1)^2^ with two basins of attraction in 0 and 1, associated with the small/big targets, respectively. The small circle represents a possible initial condition for the dynamics of *ψ*. (b) Ten simulated trajectories for *ψ*(*t*) according to *Eq. 2* with initial condition *ψ*(0) = 0.45 and noise amplitude σψ = 0.4 ms^-1^.

If we set the initial condition to *ψ*. = 0.5 and let the system evolve, the final state would be either 0 or 1 with equal probability. Shifting the initial condition towards one of the attractors results in an increased likelihood of leaning towards that same attractor, and ultimately its fixed point, i.e., the basin of attraction that was reached. For example, Figure 5b shows 10 simulated trajectories of *ψ*(*t*) where the initial condition was set to *ψ*. = 0.45. Since the initial condition is smaller than 0.5, most of the trajectories have a fixed point of 0. Nevertheless, due to the initial noise level, the fewer of them reach 1 as their final state.

The initial condition (*ψ*.) and the noise intensity (*σ_ψ_*) are interdependent. The closer an initial condition is to one of the attractors, the larger the noise is required to escape that basin of attraction. Behaviorally, the role of the initial condition is to capture the a-priori bias of choosing the smaller/bigger target. Though this is true, please note that a strong initial bias towards one of the targets does not guarantee the final decision, especially when the level of uncertainty is large. Because of this behavioral effect, we refer to the noise intensity *σ_ψ_* as *decisional uncertainty*.

The evolution of the dynamical system in Eq. 2 describes the intention of the decision-making process, at each trial *T*, of choosing the smaller/bigger target. Once a fixed point is reached, the intention is established. We call *ψ*^F^(*Ť*) the fixed point reached at trial *T*, i.e.,

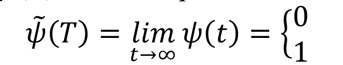

is the intended decision of choosing the smaller (0) or bigger (1) stimulus.

Although the small/big stimulus may be favored at each trial, the final decision still depends on the stimuli intensity ratio. More specifically, if the evidence associated with the small/large stimulus is higher/lower than that of its counterpart, the dynamics of the system will evolve as described in the previous section, see Eq. 1. For this reason, we incorporated the *intention* term *ψ*^F^(*Ť*) into Eq. 1, connecting the *intended decision layer* with the *neural dynamics layer*. This yields a novel set of equations

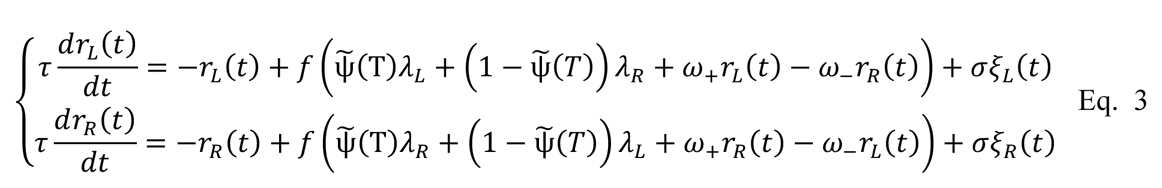

which exhibit the competence of switching preference between the large and small stimulus. If *ψ*^F^(*Ť*) = 1, the larger stimulus is favored (and the equations reduce to Eq. 1); however, if *ψ*^F^(*Ť*) = 0 the smaller is preferred.

To summarize, this *intended decision* layer endows the dynamics of decision-making hereby described with the ability of directing their preference towards either the smaller or bigger stimulus in a dynamical fashion. This inhibitory control plays the role of the regulatory criterion (size-wise) with which a decision is made in the consequential task, as described by Eq. 2.

#### 2.3.3 Layer 3: Learning the Strategy

Although the previously described intended decision layer endowed our model with the ability of targeting a specific type of stimulus at each trial, a second mechanism to internally oversee performance and to promote only beneficial strategies is a requirement. The overall goal for each participant of the consequential task is to maximize the cumulative reward value throughout an episode. As shown by previous analyses, most participants attained the optimal strategy after an exploratory phase, gradually improving their performance until the optimum is reached. Inspired by the same principle of exploration and reinforcement, we incorporated the strategy learning layer to our model.

The internal dynamics of an episode are such that selecting the small/large stimulus in a trial implies an increase/decrease of the mean value of the presented stimuli in the next trial (Figure 1). Consequently, the strategy to maximize the reward value must vary as a function of the position of the trial within episode (*T_E_*). For clarity, we labelled each trial *T* via the episode *E* and the number of trial within episode *T_E_*, i.e., *T=(E,T_E_)*. We use both notations interchangeably.

The strategy learning implemented for the model abides by the general principle of reinforcing beneficial strategies and weakening unprofitable ones, see Sec. Discussion for a comparison with existing models. At each episode *E,* the strategy function *ϕ* = *ϕ*(*Ê*, *Ť*) is updated by considering the intended choice *ψ*^F^(*Ť*) and the reward value *R(T)* obtained. In our case, this reward value originates from subjective evaluation for each individual participant in the absence of explicit performance feedback. This internal assessment yields a positive or negative perception of reward, i.e., a subjective reward. Learning implies that the preference for the selected strategy is reinforced if the subjective reward is considered beneficial. Namely, with a positive reward (*R(T)>0*), *ϕ* is increased if the larger stimulus was chosen (*ψ*^F^(*Ť*) = 1) and decreased otherwise (*ψ*^F^(*Ť*) = 0). Notice that a negative reward discourages the current strategy but promotes the exploration of alternative strategies and makes possible, eventually, to learn the optimal one over time. Mathematically, we describe the dynamics of learning as

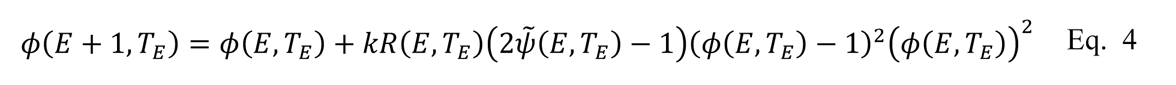

where *k* is the learning rate. Note that if *k=0*, *ϕ*(*Ê*, *Ť*) remains constant, i.e., there is no learning. The term (*ϕ*(*Ê*, *Ť*) − 1),’*ϕ*(*Ê*, *Ť*)<, is required to gradually reduce the increment to zero the closer *ϕ* gets to either zero or one, thus bounding *ϕ* in the interval [0,1]. The reward function *R(Ê*, *Ť)* represents the subjective reward. The only requirement for this function is that *R(Ê*, *Ť)* must be positive/negative if the subjective reward is considered beneficial or not. In the absence of explicit performance feedback, as is the case in the current task, participants must look for clues that convey some indirect information about their performance that could feed their internal criterion of assessment. In our case, the correct clue to look for was the change in the stimuli mean *M* between consecutive trials within an episode. For this reason, in our simulations we use *R*(*Ê*, *Ť*) = *M*(*Ê*, *Ť* + 1) − *M*(*Ê*, *Ť*) in Eq. 4. This function could be generalized in case of a different task, as discussed in the conclusions section.

Complementary to the lower layers, the strategy layer operates at a slower-pace, adaptive at a time scale of episodes. At the end of each episode, the strategy is updated by reinforcing/weakening the policy that has yielded a positive/negative reward. Mathematically, as mentioned before, this means that with a positive reward (*R(T)>0*), *ϕ* is increased if the larger stimulus was chosen (*ψ*^F^(*Ť*) = 1) and decreased otherwise (*ψ*^F^(*Ť*) = 0). In the long term, in the case that both the larger stimulus is repeatedly chosen and positive rewards obtained, then *ϕ* converges to 1. Otherwise, if both the smaller stimulus is repeatedly chosen and positive rewards obtained, then *ϕ* converges to 0. This update manifests in the next episode as a change in the initial condition for the intended decision *ψ* (Eq. 2), i.e., suggesting the direction for the intended decision to go. As shown in Figure 5, shifting the initial condition towards one of the two basins (0 or 1) increases the likelihood of reaching it. In other words, the closer the initial condition to zero/one, the more likely the intended decision will be small/big. Mathematically, this can be implemented by setting *ψ*(0) = *ϕ*(*Ť*) for each trial. In other words, the connection between the intended decision and the strategy layers lays in the influence the strategy learning exerts at each decision.

To conclude, our model consists of a three concurrent layer structure. The dynamics of each layer are defined by Eq. 3 (neural dynamics), Eq. 2 (intended decision), and Eq. 4 (strategy learning). Figure 6 shows a schematic of the model here described. The bottom part depicts the neural dynamics originated from two pools of neurons encoding the responses to two external stimuli (*L, R*). The middle (in yellow) shows the intended decision layer at every trial. Finally, the top (in green) presents the strategy learning layer, which evolves at a much slower timescale; the combined information of the intended decision and the subjective reward drives the learning of the strategy.

**Figure 6.**
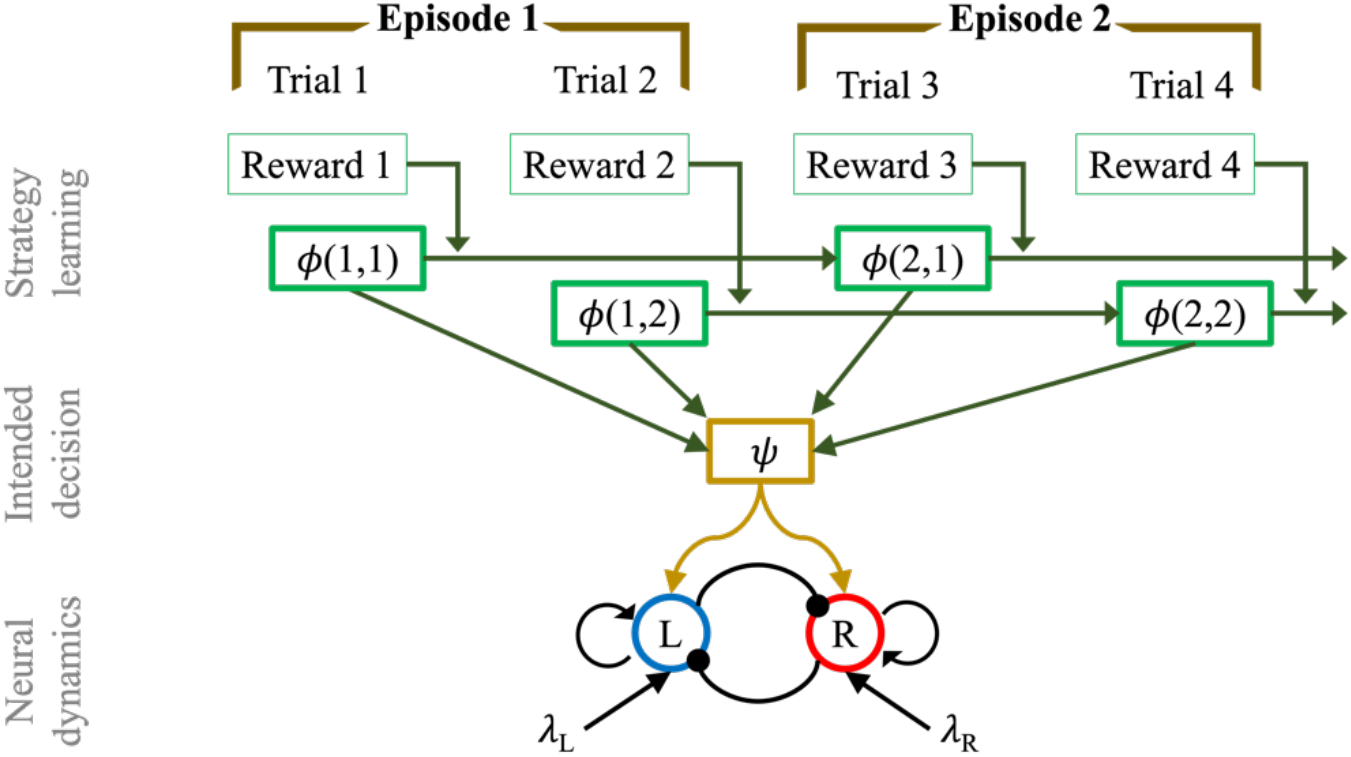
Multi-layer network structure of mean-field model of consequence-based decision making, in the case of a horizon 1 experiment. From the bottom: Neural dynamics layer: pool L is selective for the stimulus L (λL), while the other population is sensitive to the appearance of the stimulus R (λR). The two pools mutually inhibit each other (ω-) and have self-excitatory recurrent connections (ω+). The dynamics of the firing rate of the two populations is regulated by *Eq. 3*. Intended decision layer: the function ψ represents the intention, in terms of decision process, made at each trial T, of aiming for the smaller or bigger target. The dynamics of the intended decision is regulated by *Eq. 2*. Strategy learning layer: after each trial the strategy is revised, in a reinforcement learning fashion, depending on the magnitude of the gained reward value. The strategy is updated according to *Eq. 4*.

### 2.4 Model Simulations

We performed a parameter space analysis to assess the influence of the model parameters on the main behavioral metrics of interest: reaction time (RT) and performance (PF). To obtain meaningful biophysical results for the neuronal dynamics, we simulated our model varying the time constant τ, the noise amplitude σ, and the decision threshold Δ (in Eq. 3) in the following ranges: *τ* ∈ [25,95] *ms*, *k* ∈ [10^’2^, 10^’,^] *ms^-1^*, and M ∈ [0.01,0.035] *ms^-1^* (see (55)). Also, we set F_max_= 0.04 *ms^-1^*, θ = 0.015 *ms^-1^*, *k*^F^ = 0.022 *ms^-1^*, ω_+_ = 1.4, ω_-_ = 1.5. We decided to keep most of the parameters fixed (as in (55)), i.e., the ones defined within the function *f* (see Eq. 3) and the strengths of connection between pools of neurons (ω_+_ and ω_-_). As we will see below, by only varying 1, σ, and Δ we can simulate a wide range of different behaviors. In Eq. 2, we set 1*_\J_*=10 *ms* such that the dynamics of Eq. 2 is faster than the dynamics of Eq. 3 while remaining the same order of magnitude. Figure 7a shows how RT is affected by 1 and ý. By increasing the time constant τ, the RT increases both in mean and standard deviation (see FigSupp 4 a, d). The same trend occurs when increasing the threshold Δ (FigSupp 4 b, e). When varying the noise σ, we did not find a substantial difference in the RT (FigSupp 4 c, f). By fixing 1, σ, and ý, we studied the influence of the learning rate *k* and the decisional uncertainty *σ_\J_* on the PF, and, consequently, on the learning time *t_L_* (defined as in Sec. Behavioral Results). Figure 7b shows that learning time decreases as learning rate *k* increases, and as decisional uncertainty *σ_\J_* decreases. Note that for these simulations we used *n_H_=1* with 50 episodes, therefore any *t_L_* bigger than 50 means that the optimal strategy was not learned. As a consequence of this analysis, to be able to obtain a large variety of behavioral results, in the following section we vary *σ_\J_* and *k* in the following ranges: *σ*_-_ ∈ [0.2,1] *ms^-1^* and *k* ∈ [0,3].

**Figure 7.**
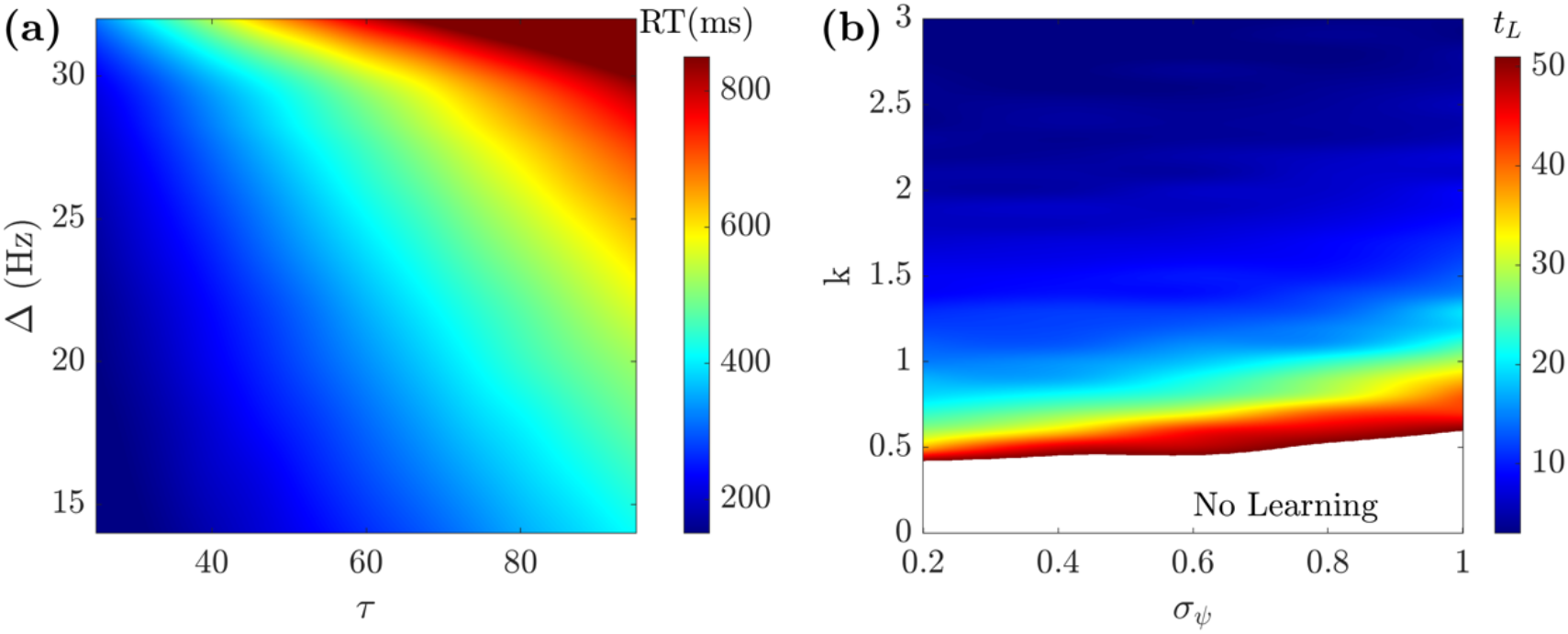
Parameter space analysis. **(a)** The RT increases when increasing either *τ* or Λ1 (σ=0.001 ms^-1^). **(b)** The learning time (tL) decreases when increasing the learning rate k and decreasing the decisional uncertainty σ_ψ_ (*τ*=81ms, σ=0.001ms^-1^, and Δ1=30 Hz).

To demonstrate the behavior of the model, Figure 8 shows the results of a typical simulation of a horizon *n_H_ =1* experiment. Figure 8a shows the example dynamics of the neural dynamics layer of our model together with the stimuli used in the simulation during the first three episodes. More specifically, the bottom row shows the time course of the two population firing rates (Eq. 3) encoding the stimuli L, R depicted in the top row. To better understand the progression of this process over time, Figure 8b gives an outlook of 36 episodes. The top row shows the performance and difficulty (in terms of difference between stimuli DbS) metrics. Note that the optimal strategy in this simulation was learned and applied from the 17^th^ episode onward. After this point, only the most difficult episodes (smallest DbS) managed to diminish the performance. The same conclusions can be drawn by looking at the middle inset, indeed after the 17^th^ episode, the intended decision metric exhibits the same pattern (small for *T_E_=1*, and big for *T_E_=2*) repeatedly. The bottom row shows the strategy learning. For the first trial within episode (*T_E_=1*), *<φ* tends to 0, i.e., it pushes the intended decision to choose the smaller stimulus. For the second trial within episode (*T_E_=2*), the trend is reversed, capturing indeed the optimal policy.

### 2.5 Individual Participants’ Behavioral Fit

This section describes the fit of the model parameters to the participants’ individual behavioral metrics. The fitting process is described as a pipeline process. In the first step, the goal is to find the best fit for the neural dynamics by fitting the reaction time (RT) and the visual discrimination (VD), i.e., fit the parameters involved in Eq. 3. We then focus on the behavioral part. The second step consists of calculating the initial preferential bias *ý_0_*. Finally, in the third step, we ran the model using the previously established parameters, and found the best fit for *σ_ψ_* and *k*, i.e., the decisional uncertainty and the learning rate. The reason why we fit the parameters in a sequential fashion is the following. The estimates of both RT and VD depend uniquely on Eq. 3. In order to evaluate the dynamics of the perceptual processes, RT and VD are fit using horizon *n_H_=0* only. Once these have been established, we focus on the behavioral part, by fitting the initial preferential bias, the learning rate and the decisional uncertainty.

**Figure 8.**
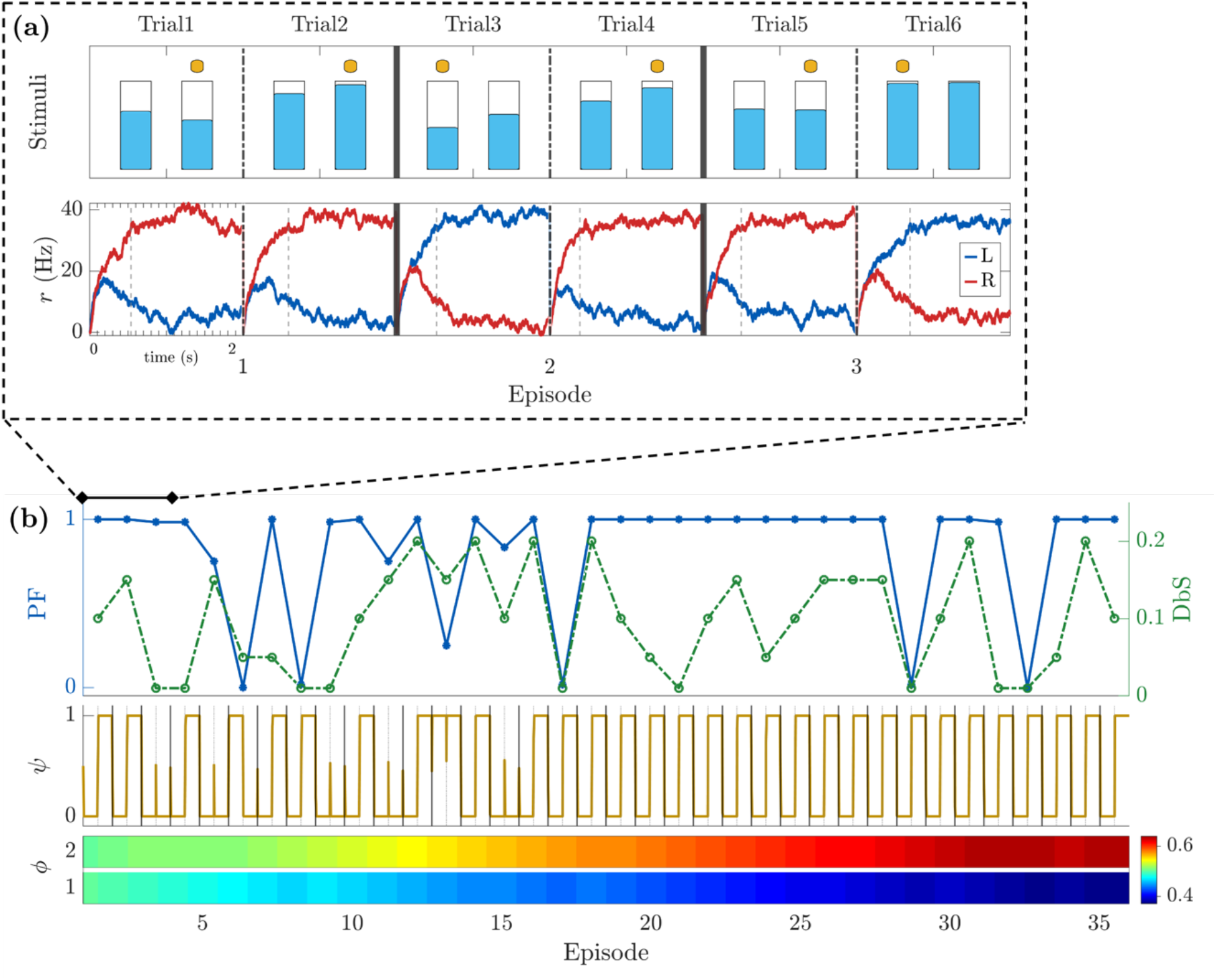
Model example simulations for a horizon 1 block. (a) Simulation of the first 3 episodes. Top row: Stimuli presentation with respective selection made in each trial displayed with a yellow dot. Bottom row: firing rate of the two populations of neurons encoding the left (in blue) and right (in red) stimuli (*Eq. 3*). Vertical dashed bars indicate the time the decision threshold was crossed. (b) Simulation of 36 consecutive episodes. First row: Performance (blue - solid) and difference between stimuli DbS (green - dashed). Second row: intended decision dynamics of choosing the bigger (1) or smaller (0) stimulus. Third row: evolution of strategy learning for each trial within episode (TE). Parameters used for the simulations: G=0.3, /-=25Hz, 1=80 ms, σ=0.006 ms^-1^, *ϕ* (1, *Ť*_E_) = 0.5 for T_E_=1,2, k=0.4, σ_ψ_=0.4 ms^-1^.

#### 2.5.1 Reaction Times and Visual Discrimination

The fitting of the model parameters to each of the participant’s behavioral metrics was performed in stages. First, we started by considering the neural dynamics layer, and fitting each parameter of Eq. 3. The first metric to fit is each participant’s RT. Note that due to response anticipation of the GO signal, the experimental RTs could be negative in a few cases (see Figure 3c). A free parameter was incorporated into the model to control for this temporal shift.

The second metric to fit is the VD, i.e., the ability to distinguish between stimuli. We assumed VD to be specific to each participant, and constant across blocks of each session. As a means of assessment, we checked how often the larger stimulus had been selected over the last 50 correct trials of the *n_H_=0* block for each level of difficulty. The only case where accuracy was low was the highest difficulty level (DbS = *0.01)*. For our model to capture this aspect, we used a linear transformation *S*° = *β* + *βs* to re-scale the stimuli *s*, ranging from 0 (empty) and 1 (full), to a range of meaningful stimuli for the model (*λ*_%,(_∼10^’,^, [22]). Furthermore, additional constraints were set for α and ý, such that this transformation did not swap the intensities between stimuli (i.e. if *S*_%_ ≥ *S*_(_ then *S*°_%_ ≥ *S*°_(_), and that the input stimuli were always positive (*S*°_%,(_ > 0). Abiding by these conditions, we varied α and ý and ran a grid-search set of simulations of Eq. 3 (with DbS |*S*_%_ − *S*_(_| = 0.01). We calculated the frequency with which the firing rate of the population encoding the larger stimulus was bigger than the alternative. The result depends not only on α and ý, but also on 1, α, and ι¢1 (see Supplementary Figure 2). Thus, to capture the large variety of results encompassed by the ranges of 1, α, and ι¢1 (see Sec. Model simulations for the respective ranges of values), while abiding by the aforementioned constraints, we let α vary between -0.03 and 0, and β vary between 0 and 0.055-2.5α. These ranges allowed for proper exploration of the parameter space.

We ran 100-trial simulations of a horizon *n_H_=0* block for each combination of the parameters 1, α, ý, and α. We then calculated the empirical cumulative distribution functions (CDF) of the RTs for all trials, and the VDs only for the difficult trials, i.e., when the DbS is *0.01*. The distribution of simulated RTs was then compared with the distributions of experimental RTs by means of the Kolmogorov-Smirnov distance (KSD) between CDFs (73–76). Since both RTs and VDs strongly depend on all parameters, both were fit simultaneously. Namely, we consider the error metric *M*^l^ = *KSD* + |^45)^ − *VD^*sim*^* |, with *c* being a constant set to 0.2 to balance the weight of the two metrics, and VD^sim^, VD^real^ being the VD from the simulated and real data, respectively. The parameters 1, α, ý, and α that minimize *M*^l^ are selected for the fit. Figure 9a depicts the CDF of the RT for the participants and for the best-fit model simulation.

To summarize, in the first step of the fit, we focused on the neural dynamics layer fit all the free parameters of Eq. 3, i.e., 1, α, ý, and α, concerned with RT and VD. The following steps will consider the behavioral component of the data.

**Figure 9.**
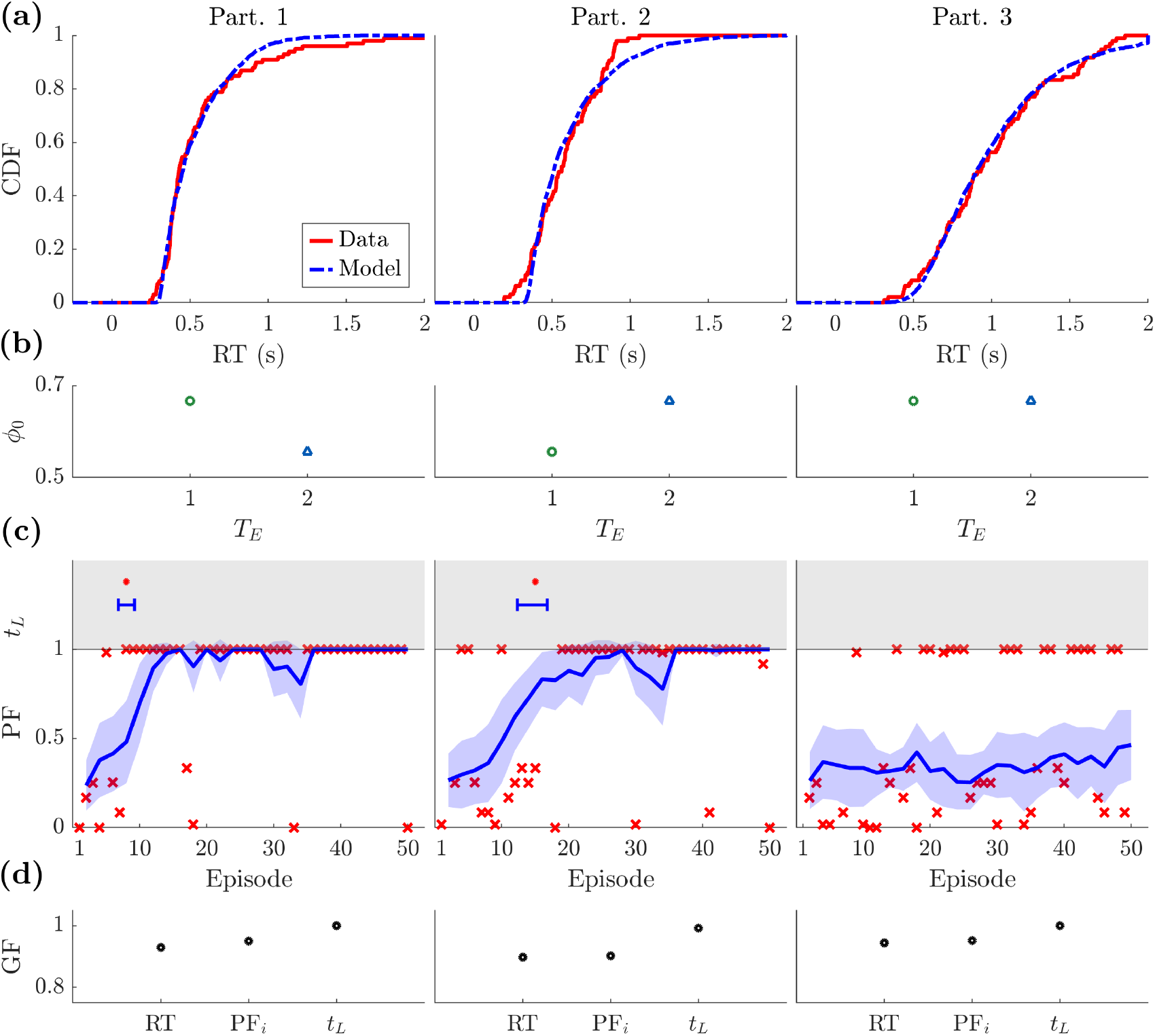
Model fit to three sample participants’ behavioral metrics. Data used: one block of horizon 1. The specific parameter values of the fit are displayed in *Table 1*. **(a)** Cumulative distribution function (CDF) of the reaction times (RT) for the participant data (solid red) and model simulation (dashed blue). **(b)** Initial bias ý0 of the participant at the beginning of the block for each trial within episode (TE). The more the preferred choice tends towards choosing the larger (smaller) stimulus, the bigger (smaller) ý0 is. **(c)** Bottom: Performance of the participant (red crosses) and of the model’s simulations (blue line: mean, shaded area: confidence interval). Top: Learning time for the participant (red dot) and model simulations (blue error bar). **(d)** Goodness of fit (GF) for three metrics: reaction time (RT), initial performance (PFi), and learning time (tL). Goodness of fit is calculated as follows: RT = 1-Kolmogorov-Smirnov distance between CDF, PFi = 1-mean square error, tL: 1-difference between learning times of participant and model’s mean divided by the total number of episodes.

#### 2.5.2 Initial Preferential Bias

Each participant performing our current task might have an initial choice preference, i.e., a natural bias towards the larger (or smaller) stimulus. In our model this is captured by the parameter *ý_0_* in Eq. 4. In the absence of bias *ý_0_* equals 0.5. The greater the preference towards the bigger choice, the closer to 1 *ý_0_* will be.

We set a vector of initial conditions *ϕ*(*Ê* = 1, *Ť*) = *ϕ*.(*Ť*) for each trial within episode (*T_E_*). To quantify *ý_0_*, we selected the first 3 episodes for each participant, and calculated the frequency *f* with which the larger stimulus was selected. The parameter *ý_0_* works as an initial condition for the intended decision process (see Eq. 2). In agreement with the attractor dynamics, if the initial condition coincides with one of the basins of attraction, the system will be locked in that state. To prevent this (since *ý_0_* should only be an initial bias), we rescaled the frequency of the selected choices *f* to make the value closer to 0.5, i.e., *ϕ*. = (1 + *f*)⁄3 (other rescaling factors could be used and would not change the results). Figure 9b shows the values obtained for *ý_0_* for each trial within episode *T_E_*. Note that we have selected one block from *n_H_=2* for participant 2 and *n_H_=1* for the others.

#### 2.5.3 Learning Rate and Decisional Uncertainty

Finally, to fit the remaining parameters **σ*_\J_* and *k* to each participant’s data, we ran the model using the previously established parameters (1, *σ*, ý, α, and *ý_0_*) and fitted its resulting performance to that of each participant. For each set of **σ*_\J_* and *k*, we ran 50 simulations and extracted the performance mean and standard deviation. To compare model and participant performances, we considered different metrics such as maximum likelihood, Bayesian (BIC) and Akaike information criterions (AIC) (74,76–79). While these are accurate methods to compare model performance, these metrics disregard the specific time dependency throughout each block, which is a key factor to characterize the learning process of the participant. In particular, the classical maximum likelihood would be strongly affected by those trials that have low performance due to errors given by fatigue or distraction. This would render this metric not suitable for our purpose. More complex methods have been recently developed to overcome this issue, such as in (80). Nevertheless, for our task we do not need such complex metrics, since our purpose is only to show that the model can fit the participants’ data and not to have a general statement on the best fit when comparing with other models. To this goal, we designed an ad-hoc novel metric consisting of two factors that determine the best fit of the learning process. The first is the initial condition, obtained by calculating the mean-square error of the performance between the model and the data during the first five episodes. By minimizing the mean-square error, we ensured that the learning process began under similar conditions for the model and for the participant. The second factor is the time required to learn the strategy. As already introduced in the Behavioral Results Section, we defined the time at which the strategy was learned as the moment after which the optimal strategy was employed in at least 9 out of the following 10 episodes, and 75% of the remaining episodes until the end of the block. To ensure that a low success rate was not due to errors caused by visual discrimination, we excluded the episodes with DbS *0.01* from this part of the fit. In summary, by combining the results for the initial conditions (*I*) and the learning time (*L*), we could extrapolate the best fit for *σ_ψ_* and *k* by minimizing the linear combination *L* + 0.1 · *I*.

Figure 9c shows the participants’ performance (red marks) as well as the associated best-fit model performance (the blue line is the mean, and the colored area is the 95% confidence interval). The top part of the plots depicts the learning time (*t_L_*) calculated for the participant (red mark) as well as for the best fit model simulations (blue error-bar). Table 1 shows the best-fit parameter values per participant.

All participants except one learned the strategy yielding maximum reward value. Specifically, participant 1 learned very fast (in 8 episodes). This was fitted by the model with the highest learning rate (*k=2.6*). Interestingly, even if participant 3 did not learn the correct strategy, the parameters obtained from the fitting process still reported a slow learning process (*k=0.2*). In addition to this, we noticed that a slightly higher learning rate was reported for participant 2, even if the strategy in this case was learned after 15 episodes only. The reason the learning rates for these two participants are similar, even though they reflect two distinct strategies, lays in the initial condition. Namely, participant 3 began the task with a stronger bias towards choosing the larger stimulus (*ϕ*.(*Ť*) = {0.67,0.67} against {0.56,0.67} for participant 2). Moreover, the noise amplitude for participant 3 is higher for both the neural dynamics *k* and the decisional uncertainty *k*_-_. When combining high noise and disadvantageous initial conditions, a weak learning rate is not enough for the strategy to be learned in a block of 50 episodes.

Figure 9d shows the goodness of fit for the two main behavioral metrics we aimed to reproduce: the reaction time (RT), and the performance, in terms of initial performance (*PF_i_*) and learning time (*t_L_*). To measure the goodness of fit, while remaining consistent with our fitting procedure, we used the following measures. For RT we calculated the KSD, for *PF_i_* we evaluated the mean-square error, and for *t_L_* we took the difference between the participant’s data and the model’s mean divided by the total number of episodes.

To summarize, we have first found the best fit for the RT and the VD by varying all the free parameters of Eq. 3, i.e., 1, α, ý, and α. Then, we calculated the subjective initial bias *ý_0_*. Finally, employing these parameters, we found the best fit for the decisional uncertainty *σ_ψ_*, and the learning rate *k*.

Finally, we show summary results for all 28 participants. To illustrate that the model is able to capture all participants’ behavioral results, Figure 10 shows the goodness of fit for the RT, initial performance PF_i_, and learning time t_L_ for the entire set of 28 participants. For all three metrics, we show the scatter plot including each participant, the respective distribution, and the boxplot depicting the median and the 25th/75th percentile. For reference, we superposed colored markers on the results of the three sample participants shown in the previous figure.

**Figure 10.**
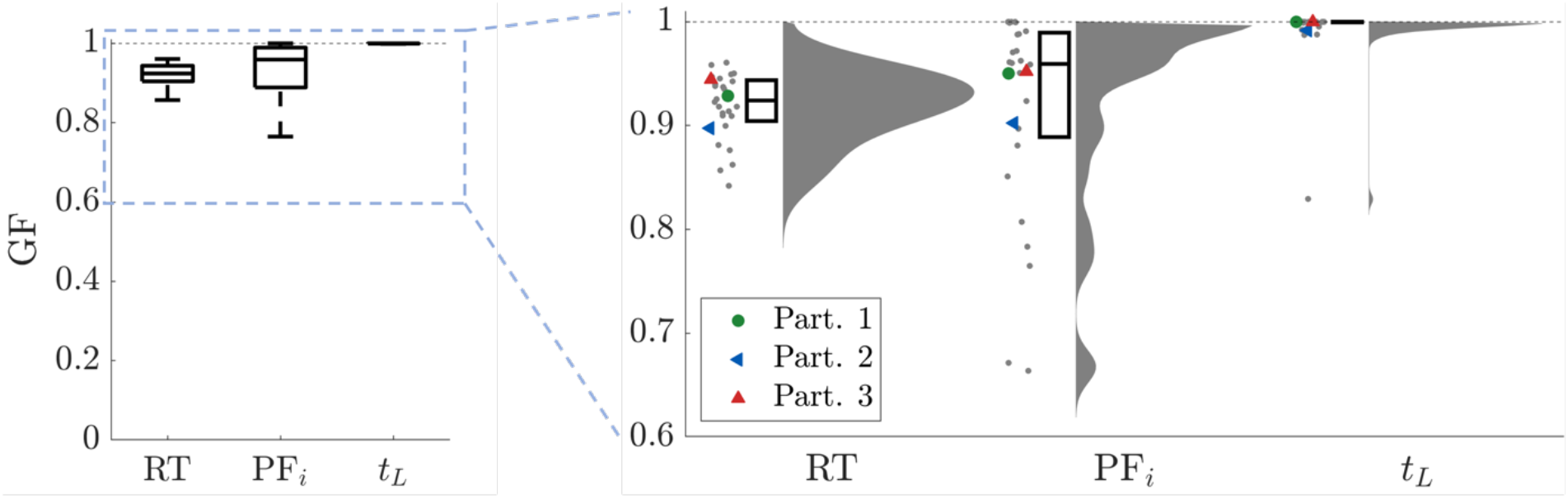
Goodness of fit. For RT we calculated KSD, for PFi we evaluated the mean-square error, and for tL we took the difference between the participant’s data and the model’s mean divided by the total number of episodes. For all three metrics, we show the scatter plot of each single participant, the respective distribution, and the boxplot depicting the median and the 25/75 percentile. For reference, we superposed (colored markers) the results for the three participants shown in the previous figure.

To summarize, we performed an individual fit to each of the participant’s behavioral metrics. We first used the RT distribution and VD of each participant to fit the parameters in Eq. 3. Once these parameters were fixed, we moved on to calculate the initial bias, and ran simulations of the model. Finally, we compared the results of the simulations with the performance of the participants and found the best fit for the behavioral parameters, i.e., the learning rate and decisional uncertainty.

## 3 DISCUSSION

Here we provided a characterization of long-term consequence-based option assessment during decision-making, and a plausible theoretical account of its underlying neural processes. To this end, we designed an experimental task in which trials were grouped into episodes of one to three trials and choice influenced the reward value of stimuli in subsequent trials. The stimuli shown in trials within each episode were deliberately designed to promote inhibitory choices first and an incentive one in the last trial. To specifically characterize how a consequence-based assessment forms and influences decisions as a function of learning, we instructed each participant to explore his/her decisions to find the strategy yielding the most of cumulative reward value within episode in the absence of any explicit performance feedback. Our purpose was to promote the participant to develop his/her own subjective assessment of performance, based on relating the size of the stimuli in the trial next to the choice in the previous trial. Remarkably, most participants attained the optimal strategy. This demonstrates that they grasped the relationship between their decisions and the consequence, incorporating their predictions of future choice options to their internal assessment of performance, and biasing their policy consistently with a maximization of cumulative reward value.

Some similarities can be found when comparing our task with the *farming on Mars* task (53). In this task participants were asked to make repeated choices between two alternatives with the goal of maximizing the rewards they receive over the entire session. Each time a participant selects the more attractive alternative, the future utility of both options is lowered. Essentially, the *farming on Mars* task would be the equivalent of 1 episode of horizon 100 of our consequential task. We claim that our task is an extension of the farming on Mars task, and it is more appropriate when studying different aspects of consequence. First, having more repetitions, of shorter horizon, made the learning process different. Namely, limiting the horizon means reducing the time available to unravel the optimal strategy and it promotes the generalization to different horizon depths. Furthermore, using such a structure allows us to study the impact of consequence on the different trials within episodes *T_E_*, i.e., with or without consequence. For instance, the RT is faster for the last *T_E_* because there is no consequence. Moreover, our task is more flexible since the optimal policy, to maximize reward, can be easily changed (for example, big-small-big for *n_H_* 2) to study how participants would adapt to the change of strategy. In addition, by changing only one parameter value, our task can be modified such that the optimal policy becomes stochastic. Namely, by decreasing the gain/loss parameter *G* (see Materials and Methods - episode structure), the maximum cumulative reward could be attained by always choosing the bigger stimulus, when the difference between stimuli is large enough to compensate the loss *G* across trials. Finally, our task was designed to be performed by humans and, with only small changes, non-human primates. This opens a new set of possible analyses that can be done, such as studying the neural dynamics for different *T_E_* and horizon depths.

In addition to the experimental analyses, we introduced a novel mathematical model encompassing the cognitive processes required for consequence-based decision-making in a joint framework. The model is organized in three layers. The bottom layer describes the average dynamics of two neural populations, representing each the preference for one option, competing against each other until their difference in activity crosses a threshold. The middle layer implements the participant’s preference for choosing the bigger or smaller stimulus at each specific trial (the so-called intended decision). The top layer describes the strategy learning process, which oversees the model’s performance, adapts by reinforcement to maximize the cumulative reward value, and drives the intended decision layer. This oversight mechanism, combined with the modulation of preference, accurately reproduces an internal process of consequence assessment and subsequent policy update. The model was validated by fitting its parameters to reproduce each participant’s behavioral data (reaction time distribution, visual discrimination, initial bias, and performance). The model predictions faithfully reproduced these metrics along with the learning time for each participant, regardless of their level of accuracy throughout the session. Importantly, this model also provides a plausible account of the neural processes required for option gauging as a function of their associated consequence in terms of reward, and of how these processes participate in decision-making.

### 3.1 Rule-Based vs Far-Sighted Assessment of Consequence

The optimal strategy to attain maximum cumulative reward value may be operationalized as a sequence of decision rules: choose small, then big in horizon 1 episodes; choose small, then small, then big, in horizon 2 episodes. Although we expected the participants’ choices to abide by these rules once the learning was complete and the optimal decision strategy established, the focus of this study is on how consequence-based assessment forms and influences the learning of decision-making strategies. Because of this, it was crucial to run a task design devoid of any explicit performance feedback, which could potentially inform the participant of his/her performance throughout each episode and ultimately promote a rule-based strategy from the very beginning.

For the same purpose, and to promote exploration, the participants were left in the uncertainty of neither having a clear criterion to decide upon nor the knowledge about which aspect of the stimuli to prioritize to obtain bigger reward values in the trial next and across the episode. Note that, in addition to the height of the bars (proportional to reward value), the stimuli at each trial were presented on the right and left of the screen, and were shown sequentially, randomly alternating their order of presentation across trials. Although meaningless from the perspective of gaining the most of reward value, both the position and order of presentation contributed to increase the uncertainty as to which dimension of the stimuli were relevant to attain the goal during the learning phase. In fact, under these conditions, the participants were left with a single element that could aid them build their internal criterion to assess performance: perceiving the relationship between their choice at a trial, and the stimuli being subsequently presented in the next. If noticed, over a few episodes, this piece of evidence could then be used to predict the consequence associated with choosing each option at each trial within episode. To this end, participants had to rely on their own subjective perception of performance, fed alone by their observations of the stimuli presented after each decision, and by their own internal assessment criterion, based on their skill at estimating the sum of water (reward value) throughout the trials of each episode. Importantly, learning the optimal strategy could only be achieved via exploration, either purposely or randomly, testing the pairing between the stimuli presented at each trial, the choice made, and, most importantly, the stimuli of the trial next.

To summarize, the problem of having explicit performance feedback is that the learning of the optimal strategy could be reduced to testing rule-based sequences until the one that gives the optimal feedback is found. Although the optimal strategy consists of the same rule-based sequence, the crucial element of the task is that, to reach that stage, the participant must first forego a phase of exploration in which learning is driven by exploration and assessment of the reward-based consequence associated with each option. Until then, the learning depends on a computation of reward value encompassing the consideration of far-sighted effect of each decision within episode, on the grounds of an internal subjective assessment criterion that makes this learning possible, and the results hereby presented non-trivial.

### 3.2 Building a Subjective Assessment Criterion

The crucial element of the aforementioned process is that, in the absence of explicit performance feedback, learning depends on first building up an internal criterion of reward. This criterion necessarily depends on cognitive processes implementing an oversight mechanism of whether the correct decision criterion is being used, and whether the proper association between the choice and subsequent stimuli is being correctly perceived (81–84). Moreover, despite the participants being able to find the optimal strategy and diminishing the uncertainty of their behavior to reach the optimal strategy, the fact they never get an explicit external confirmation forces them to bear the doubt of whether their strategy is indeed the optimal one. The discussion of the theoretical formalization presented next suggests a minimal implementation for these mechanisms. This suggests a plausible strategy for this subjective mechanism to capture the relationship between stimuli and subsequent stimuli are established on a single trial basis, within the wider decision-making strategy of maximizing cumulative reward value.

### 3.3 Computational models of consequence

The analyses described in the results section demonstrate that the consequential task is an appropriate framework to study how consequence-based option assessment forms and influences decision-making. In parallel, the model we developed has the goal of reaching a formal characterization of the cognitive processes underlying the operations necessary to perform this task. As for most value-based decision-making models (23,69,85–88), learning in our model is operationalized by a reinforcement comparison algorithm, scaled by the difference between predicted vs. obtained reward value (89,90), measured accordingly to the participant’s subjectively perceived scale. For simplicity, we assumed a fixed function across participants to quantify reward value (R(T) function in Eq. 4). Furthermore, to provide the necessary flexibility for the model to capture the full range of participants’ learning dynamics, the model included two free parameters, the learning rate and the decisional uncertainty, to be fit to each participant’s behavior. The result is a model that could faithfully reproduce the full range of behaviors of each participant: RT distribution, pattern of decision-making, and learning time.

The structure of the model is organized in three layers. The lower layer (neural dynamics) reproduces the average activity of two neural populations competing for the selection, each representing one of the two stimuli to decide upon at each trial. The commitment for one of the two options is taken when the difference in firing rate between the two populations crosses a given threshold (23,55,85). A similar architecture, with small variations, has been used to model decision-making in a broad set of tasks (21,55,91,92) and can describe most types of single-trial, binary decision-making, including value-based and perceptual paradigms. Beyond the scope of this investigation, this model can also subserve working memory (21,93); a transient input can bring the system from the resting state to one of the two stimulus-selective persistent activity states, which can be internally maintained across a delay period. However, modelling consequence-based decision-making requires at least two additional mechanisms beyond binary population competition. The first is to surmise criteria to prioritize a specific policy for decision-making. The second is to create an internal mechanism of performance to evaluate these criteria, based on the difference between predicted and obtained reward value. Accordingly, the role of the middle layer (intended decision) is the implementation of those criteria, which in our case depends on the relative value of the stimuli and on the number of trial within episode. Finally, the top layer (strategy learning) implements the learning by reinforcement comparison (94) and temporal difference (89,95).

We claim that the model is a minimal implementation of consequence-based decision-making within the context of our experimental task. Each part of the model is in fact essential to describe decision-making, inhibition, and learning. For the neural dynamics layer, the set of equations corresponds to most reduced version of a network of spiking neurons for binary decision-making (21); it makes use of only 2 populations of neurons and a minimal set of parameters. The middle layer consists of one equation (with only one free parameter) and makes use of the simplest possible shape for the description of two-attractor dynamical system with the addition of a noise component. Finally, the top layer follows the same type of implementation as a classical temporal difference algorithm, and only adds one free parameter to the model, i.e., the learning rate. Each of these three layers is indispensable for a biologically plausible theoretical formalization of consequence-based decision-making. Indeed, without the first layer we would not have a biologically plausible decision-making, without the middle layer we could not describe the change of policy, and without the top layer we would not have learning.

In the existing literature, there are models that describe learning processes during decision-making. One of the most popular classes of such models is reinforcement learning (RL) (94). Among RL models, temporal difference algorithms, such as Q-learning, are often used to model behavior. To use these types of models, one must define the state-action space representing a particular context. For *n_H_ 1* of the consequential task, for example, one could define a 3×2 Q-table where the action space consists of choosing either the smaller or the larger stimulus and the state space represents *T_E_ 1, T_E_ 2_low,* and *T_E_ 2_high*. From there, an epsilon-greedy Q-learning algorithm could be used to learn the optimal strategy by continuously updating the estimated value associated with each state-action pair upon visiting them. The closest model to the one we built here is the RL drift diffusion model (96). This model can reproduce both RT and choices patterns. The advantage of our model is that it not only describes behavioral patterns of learning, but it is also biophysically plausible when describing RT. Moreover, since the neural dynamics of the mean-field approximation have been derived analytically from networks of spiking neurons (61), a direct link from neural data and this model could theoretically be achieved.

The results and predictions depicted in the model descriptive section show that the dynamics of the three layers combined can accurately reproduce the behavior of each single participant, including those who did not attain the optimal strategy. The low number (4) of equations in the model, together with the low number (7) of free parameters, makes this model a simple, yet powerful tool able to reproduce a large variety of behavioral results. Moreover, unlike the basic reinforcement learning agents or models for evidence accumulation, our model is biologically plausible and therefore able to fit individual behavioral metrics, such as RT, initial bias, and visual discrimination. Note that, for the behavioral part of the model, the free parameters are only 3, i.e., learning rate, decisional uncertainty, and initial bias. The same number of free parameters is needed for classical reinforcement learning algorithms, e.g., Q-learning.

The comprehensive formulation of the model makes it possible to explain and fit various scenarios. We have already mentioned the differences in learning speeds, and that the model could fit any of them, even when there is no learning. Another example is the difference in the order of execution of the blocks. Namely, most participants when they learned the optimal strategy in one horizon, they generalized the rule and applied it to the other horizon block, making the learning much faster (see Supplementary Materials). In our model, this is captured mainly by the initial bias, that is calculated for each block individually. As third example, potentially, a characteristic that our model could fit is the difference in RT between trials within episodes and horizons (see Figure 2f). In this manuscript, for simplicity, we decided to perform a single fit for the neural dynamics’ equations, finding one set of parameters per participants. To explain the differences between horizons and trials within episodes, the same fit should be done for each condition. Moreover, even if it is not the case of this specific task, the model is able to adapt in case of a sudden change of strategy. Nevertheless, if this would be the case, it would be advisable to adopt a more realistic adaptation mechanism. Namely, it seems reasonable to assume that, after learning, once a participant realizes that the optimal strategy used so far is not working anymore, he would reset his strategy instead of gradually change it. However, even though it is an interesting topic, this is work for future investigation.

## 4 Conclusion and Future Work

In this manuscript we have introduced a novel minimalistic formalism of the brain dynamics of consequence-based decision-making and its associated learning process. We validated this formalism with the behavioral data gathered from twenty-eight human participants, which the model could accurately reproduce. By extension of the classic single-trial binary decision-making, we designed a mechanism of oversight based on the assessment of the effect of previous decisions on subsequent stimuli, and a reinforcement rule to modify behavioral preferences. As part of the same study, we also designed the consequential task, a novel experimental framework in which gaining the most of reward value required learning to assess the consequence associated with each option during the decision-making process. Both the experimental results and the model predictions review consequence-based decision-making as an extended version of value-based decision-making in which the computation of predicted reward value may extend over several trials. The formalism introduces the necessary notions of oversight of the current strategy and of adaptive reinforcement, as the minimal requirements to learn consequence-based decision-making.

Although our model has been designed and tested in the consequential task described here, we argue that its generalization to similar paradigms in which optimal decisions require assessing the consequence associated to the options presented, or sequences of multiple decisions, may be relatively straightforward. Specifically, we envision three possible future extensions to facilitate its generalization. First, the model could incorporate several preference criteria simultaneously or combinations thereof to the intended decision layer: left vs. right or first vs. second, instead of small vs. big, to be determined in a dynamical fashion. This could be achieved with a multi-dimensional attractor model, with as many basins of attraction as the number of preference criteria to be considered.

The second future extension is the re-definition of the reward function R(T) according to the subjective criterion of preference. Namely, a reward value can be perceived differently by different participants, i.e., people operate optimally according to their own subjective perception of the reward value. Because of this, a possible extension is to incorporate an individual reward value function per participant (R(T) in Eq. 4). For simplicity, in this manuscript we set R(T) to be fixed and to be the objective reward value function. In case a participant did not perceive what was the optimal reward value, he/she performed sub-optimally according to objective reward function, and the model responded by allowing the learning constant *k* to be zero. This holds since the optimal strategy was never reached, and the fitting of the participant’s performance was correct. Nevertheless, it remains a standing work of significant interest to investigate different subjective reward mechanisms and their implementation in the model.

Finally, the third future point to investigate is whether the learning rate is time dependent, i.e., *k(E)*. This would facilitate reproducing learning processes starting at different times throughout the session. For example, it is possible that participants initiate the session having in mind a possible (incorrect) strategy and they stick to it without looking for clues, and therefore without learning the optimal policy. Nevertheless, after many trials they may change their mind and begin to explore different strategies. In this case, the learning rate *k(E)* would be set to zero for all the initial trials when indeed there is no learning.

Again, we want to emphasize that even if this model is built for the consequential task, it contains all the elements and processes to reproduce other tasks of sequential consequence-based decision-making. Note that the strategy learning mechanism is already general enough to adapt to tasks where the optimal policy is not fixed throughout the experiment. Indeed, if the optimal policy would change suddenly at some point during the block, the learning mechanism would be able to detect a change and adapt accordingly. Finally, we want to stress that our model could be applied to other decision-making paradigms, such as a version of the consequential random-dot task (97) or other multiple-option paradigms.

## 5 MATERIALS AND METHODS

### 5.1 Participants

A total of 28 participants (15 males, 13 females; age range 18-30 years; all right hand dominant) participated in the experimental task. All participants were neurologically healthy, had normal or corrected to normal vision, were naive as to the purpose of the study, and gave informed consent before participating. The study was approved by the local Clinical Research Ethics Committee (CEIm Ref. #2021/9743/I) and was conducted in accordance with relevant guidelines and regulations. Participants were paid a €10 show-up fee.

### 5.2 Experimental Setup

Participants were situated in the laboratory room at the Facultat de Matemàtiques i Informàtica, Universitat de Barcelona, where the task was performed. The participants were seated in a chair, facing the experimental table, with their chest approximately 10cm from the table edge and their right arm resting on its surface. The table defined the plane where reaching movements were to be performed by sliding a light computer mouse (Logitech Inc). On the table, approximately 60cm away from the participant’s sitting position, we placed a vertically-oriented, 24” Acer G245HQ computer screen (1920×1080). This monitor was connected to an Intel i5 (3.20GHz, 64-bit OS, 8 GB RAM) portable computer that ran custom-made scripts, programmed in MATLAB with the help of the MonkeyLogic toolbox, to control task flow (NIMH MonkeyLogic, NIH, USA; https://monkeylogic.nimh.nih.gov). The screen was used to show the stimuli at each trial and the position of the mouse in real time.

As part of the experiment, the participants had to respond by performing overt movements with their arm along the table plane while holding the computer mouse. Their movements were recorded with a Mouse (Logitech, Inc), sampled at 1 kHz, which we used to track hand position. Given that the monitor was placed upright on the table and movements were performed on the table plane (horizontally, approximately from the center of the table to the left or right target side), the plane of movement was perpendicular to that of the screen, where the stimuli and finger trajectories were presented. Data analyses were performed with custom-built MATLAB scripts (The Mathworks, Natick, MA), licensed to the Universitat de Barcelona.

### 5.3 Consequential Decision-Making Task

This section describes the consequential decision-making task, designed to assess the role of consequence on decision-making while promoting prefrontal inhibitory control (98). Since consequence depends on a predictive evaluation of future contexts, we designed a task in which trials were grouped together into episodes (groups of one, two or three consecutive trials), establishing the horizon of consequence for the decision-making problem within that block of trials.

The number of trials per episode equals the horizon *n_H_* plus 1. In brief, within an episode, a decision in the initial trial influences the stimuli to be shown in the next trial(s) in a specific fashion, unbeknown to our participants. Although a reward value is gained by selecting one of the stimuli presented in each trial, the goal is not to gain the largest amount as possible per trial, but rather per episode.

Each participant performed 100 episodes for each horizon *n_H_* = 0, 1, and 2. In the interest of comparing results, we have generated a list of stimuli for each *n_H_* and used it for all participants. To avoid fatigue and keep the participants focused, we divided the experiment into 6 blocks, to be performed on the same day, each consisting of approximately 100 trials. More specifically, there was 1 block of *n_H_=0* with 100 trials, 2 blocks of *n_H_=1* each with 100 trials, and 3 blocks of *n_H_=2* with two of them of 105 trials and one of 90. Finally, we have randomized the order in which participants performed the horizons.

Figure 1 shows the timeline of one horizon 1 episode (2 consecutive trials). The episode consists of two dependent trials. At the beginning of the trial, the participant was required to move the cursor onto a central target. After a fixation time (500 ms), the two target boxes were shown one after the other (for 500 ms each) to the left and right of the screen, in a random order. Targets were rectangles filled in blue by a percentage corresponding to the reward value associated with each stimulus (analogous to water containers). Next, both targets were presented together. This served as the GO signal for the participant to choose one of them (within an interval of 4s). Participants had to report their choice by making a reaching movement with the computer mouse from the central target to the target of their choice (right or left container). If the participant did not make a choice within 4 s, the trial was marked as an error trial. Once one of the targets had been reached for and the participant had held that position (500ms), the selection was recorded, and a yellow dot appeared above the selected target, indicating successful selection and reward value acquisition. In case of horizons larger than 0, the second trial started following the same pattern, although with a set of stimuli that depended on the previous decision (see next section). A progress bar at the bottom of the screen indicates the current trial within the episode (for horizon 1, 50% during the first trial, 100% during the second trial).

At the beginning of the session, participants were given instructions on how to perform the task. Specifically, using some sample trials, we demonstrated them how to select a stimulus by moving the mouse. Step by step we showed that a target appears in the center of the screen indicating the start of an episode. We told them that they had 4 seconds to move the cursor to the central cross. After moving the cursor to the central cross, two bars appear, one after the other, and once both appear together/simultaneously, they had 4 seconds to make their decision by moving the cursor over one of the two bars. At that point a yellow dot appears over the bar indicating their selection. After that, the central target appears again indicating the beginning of a new trial. After explaining how to technically execute the task, we focused on explaining the task goal. We showed them a schematic of the task, much like the one in Figure 1a illustrating the structure of trials and episodes. We told them that the goal is to get as much reward (water) as possible in each episode, and that for episodes with more than 1 trial each, the choice in a trial may have an effect on what appears in the next trial in the same episode. We encouraged them to explore in order to try to figure out what that effect might be, while keeping in mind that their goal is always to maximize the total reward in each episode. Finally, we told them that they will be presented with a series of episodes in a row, each episode is independent, meaning that their decisions in one episode have no effect on subsequent ones.

### 5.4 Episode Structure

The participants were instructed to maximize the cumulative reward value throughout each episode, namely the sum of water contained by the selected targets across the trials of the episode. If trials within an episode were independent, the optimal choice would be to always choose the largest stimulus. Since one of the major goals of our study was to investigate delayed consequence assessment involving adaptive choices, we deliberately created dependent trial contexts in which making incentive decisions (selecting the larger stimulus) would not necessarily lead to the most cumulative reward value within episode.

To promote inhibitory choices, the inter-trial relationship was designed such that selecting the small (large) stimulus in a trial, yielded an increase (decrease) in the mean value of the options presented in the next trial. As explained below, because of the parameters choice we made, always choosing the larger stimulus did not maximize cumulative reward value for *n_H_=1, 2*.

Trials were generated according to 3 parameters: horizon’s depth *n_H_*, perceptual discrimination (in terms of difference *d* between the stimuli), and the gain/loss *G* in mean size of stimuli for successive trials. The stimuli *S*_9,,_ presented on the screen could take values ranging from 0 to Trials were divided into five difficulty levels by setting the difference between stimuli (DbS) *d* ∈ {0.01,0.05,0.1,0.15,0.2}.

For horizon *n_H_=0*, for each trial the stimuli *S*_9,,_ are generated as to have mean *M* and difference *d* between them, i.e., *S*_9,,_ = *M* ± *d*/2. To have stimuli ranging from 0 to 1, the mean *M* is randomly generated using a uniform distribution with bounds [*d*_)*+_/2,1 − *d*_)*+_/2], where *d*_)*+_ = 0.2 is the maximum DbS. In horizon *n_H_=1*, each episode consists of 2 dependent trials. Specifically, the stimuli presented in the second trial depend on the selection reported in the previous trial of that same episode. More specifically, the rule is such that if the choice of the first trial is the smaller/larger stimulus, the mean of the pair of stimuli in the second trial will be increased/decreased by a specific gain *G*. In practice, the first trial of an *n_H_=1* episode is generated in the same way as for horizon *n_H_=0*, i.e., the two stimuli equal *S*_9,,_ = *M* ± *d*/2. The stimuli in the second trial within the same episode could be either *S*_9,,_ = *M* + *d* ± *d*/2 or *S*_9,,_ = *M* − *d* ± *d*/2, depending on the previous decision. Note that the difficulty of the trial remains constant within episode. A schematic for the trial structure is shown in Figure 1. Again, to have stimuli ranging from 0 to 1, the mean *M* is randomly generated using a uniform distribution with bounds [*d* + *d*_)*+_/2,1 − *d* − *d*_)*+_/2]. In horizon *n_H_=2*, episodes consist of three trials. The trial generation is structured as for horizon *n_H_=1*. Namely, the first trial has stimuli *S*_9,,_ = *M* ± *d*/2, the second *S*_9,,_ = *M* ± *d* ± *d*/2, and the third *S*_9,,_ = *M* ± *d* ± *d* ± *d*/2. To have stimuli ranging from 0 to 1, the mean *M* is randomly generated from a uniform distribution with bounds [2*d* + *d*_)*+_/2,1 − 2*d* − *d*_)*+_/2]. We set the gain/loss parameter to *G=0.3* and *G=0.19* for horizon *n_H_=1* and *n_H_* = 2, respectively. Our choice was motivated by the fact that G should be big enough to have a deterministic optimal strategy, i.e., always choosing the smaller reward value apart from the last trial within episode. In other words, choosing the bigger stimulus never compensates for the loss given by G. Moreover, *G* should be big enough to let the participants perceive the gain/loss between trials, while simultaneously allowing some variability for the randomly generated means *M*.

## 5.5 Statistical analysis

We were interested in testing the relationship of the performance (PF) and the reaction time (RT) with the horizon *n_H_*, trial within episode *T_E_*, and episode *E*. To obtain consistent results, we adjusted these variables as follows. The trial within episode was reversed, from last to first, because the optimal choice for the last *T_E_* (large) was the same regardless of the horizon number. The variable representing the trial within episode counted backwards was denoted as *Ť*. Furthermore, regarding the model for PF, to consider trials within episode independently, we adapted the notion of PF (defined as a summary measure per episode) to an equivalent of PF per trial, i.e., the percentage of optimal choices *P*_*oc*_. To be able to calculate such percentage, we grouped the episodes in blocks of 10 and used their average. This new variable was called *Ê*. Regarding the model for RT, since we considered each episode separately, and not an aggregate of 10 of them, we also checked the dependency with DbS (*d*). Finally, to assess the difference between learning groups, we introduced the categorical variable L that identifies the group of participants that learned the optimal strategy and the ones who did not according to Figure 2a. We then used a linear mixed effects model (59,60) to predict PF and RT. The independent variables for the fixed effects are horizon *n_H_*, trial within episode *Ť*! (counted backwards), and the passage of time expressed as groups of 10 episodes *Ê* each for PF, or for RT the episode *E* and DbS *d*. We set the random effects for the intercept and the episodes grouped by participant *p^*; we wrote the random effects as ’*Ê*(*p^*). The resulting models are: *P*_*oc*_∼*L* ⋅ *Ê* + *L* ⋅ *n*_$_ ⋅ *Ť* + ’*Ê*(*p^*) and *RT*∼*L* ⋅ *Ê* + *L* ⋅ *d* + *L* ⋅ *n*_$_ ⋅ *Ť* + (*Ê*|*p^*). The RT, *P*_*oc*_, *Ê*, *E*, and DbS were z-scored to run the analysis. The results of the statistical analysis are reported in Table 2. The regression coefficients, with their respective group significance, are shown in Figure 2e-f.

## Acknowledgments

This project has received funding from the European Union’s Horizon 2020 Framework Programme for Research and Innovation under the Specific Grant Agreement N. 945539 COREDEM (Human Brain Project SGA3).

## Supplementary Materials

### Exploratory strategy

We analyzed the exploratory strategy participants used. In particular, we tested whether participants only considered the size of the stimuli (small/big), or if they also tried other hypotheses, such as the order of presentation (first/second) or the location (left/right) of the stimuli. The result of this analysis is shown in FigSupp 1, which depicts the proportion of trials for three exploratory strategies, i.e., size, location, and order of presentation.

Participants mostly considered the size as a possible factor for optimization during the trials when the optimal strategy was not yet found, FigSupp 1a. These panels show that some participants explored the location and the order of appearance. Nevertheless, according to a questionnaire we ran at the end of each session, asking participants what strategy they learned and/or tried, only 4 mentioned the order of appearance, and none the location.

From the 6 participants who did not learn the optimal strategy (FigSupp 1b), only 1 considered the order of appearance throughout the whole session. This participant reported in the questionnaire that they used a complex combination of appearance and size as metric for optimization. Another participant reported in the questionnaire that they performed at random for *n_H_=0,* and for *n_H_=1,2* tried to make the cumulative reward fill to the top of the bar, resulting in choosing small-big for *n_H_=1,* and small-small-big for *n_H_=2*. All other 4 participants repeatedly chose the larger stimulus for all trials.

**FigSupp 1.**
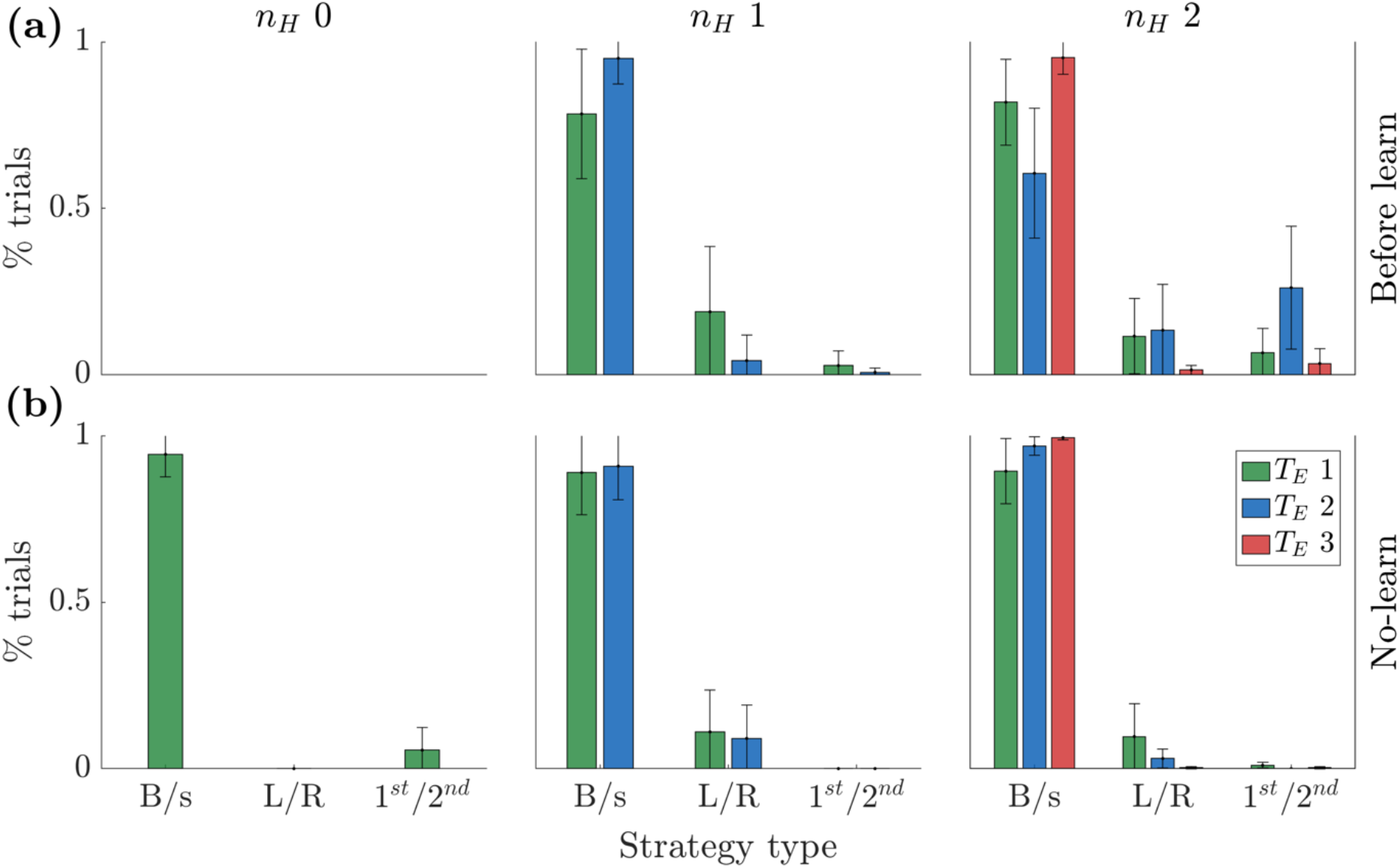
Exploratory strategy during the episodes before learning and for the participants who did not learn the optimal strategy. **(a)** Participants mostly considered the size as a possible factor for optimization during the trials when the optimal strategy was not yet found. For nH 0 there are not enough trials before learning to perform this analysis. **(b)** From the 6 participants who did not learn the optimal strategy only one considered the order of appearance throughout the whole session.

### Learning time and order of performance of different horizons

In Materials and Methods, Sec. Consequential Decision-Making task, we described the structure of the task, and in particular we mentioned that we randomized the order in which participants performed the horizons. This means that, for example, some participants performed *n_H_* 2 before *n_H_* 0. We wondered if the order of the horizons had an influence on the learning time. For example, were the participants who started with n_H_ 1 faster in learning the optimal strategy than the ones who started with n_H_ 2? To address this, we performed a thorough investigation analyzing the learning time for each block of recording of the session.

FigSupp 2 shows the learning time for each block in order of execution (x-axis). Each plot is a different participant. On the y-axis, the learning time is reported in terms of episodes, and above 50 means no-learning. Different horizons are depicted with different markers and colors: *n_H_* 0 green circle, *n_H_* 1 blue triangle, *n_H_* 2 red cross. This figure also shows that 2 participants did not learn any horizon, 4 participants learned *n_H_* 0, but not either *n_H_* 1 or *n_H_* 2, and the remaining 22 managed to learn at least one of *n_H_ 1, 2*. Note that participants 1, 2, and 3 in the main text of the manuscript correspond to participants 20, 6, and 7 in FigSupp 2, respectively.

**FigSupp 2.**
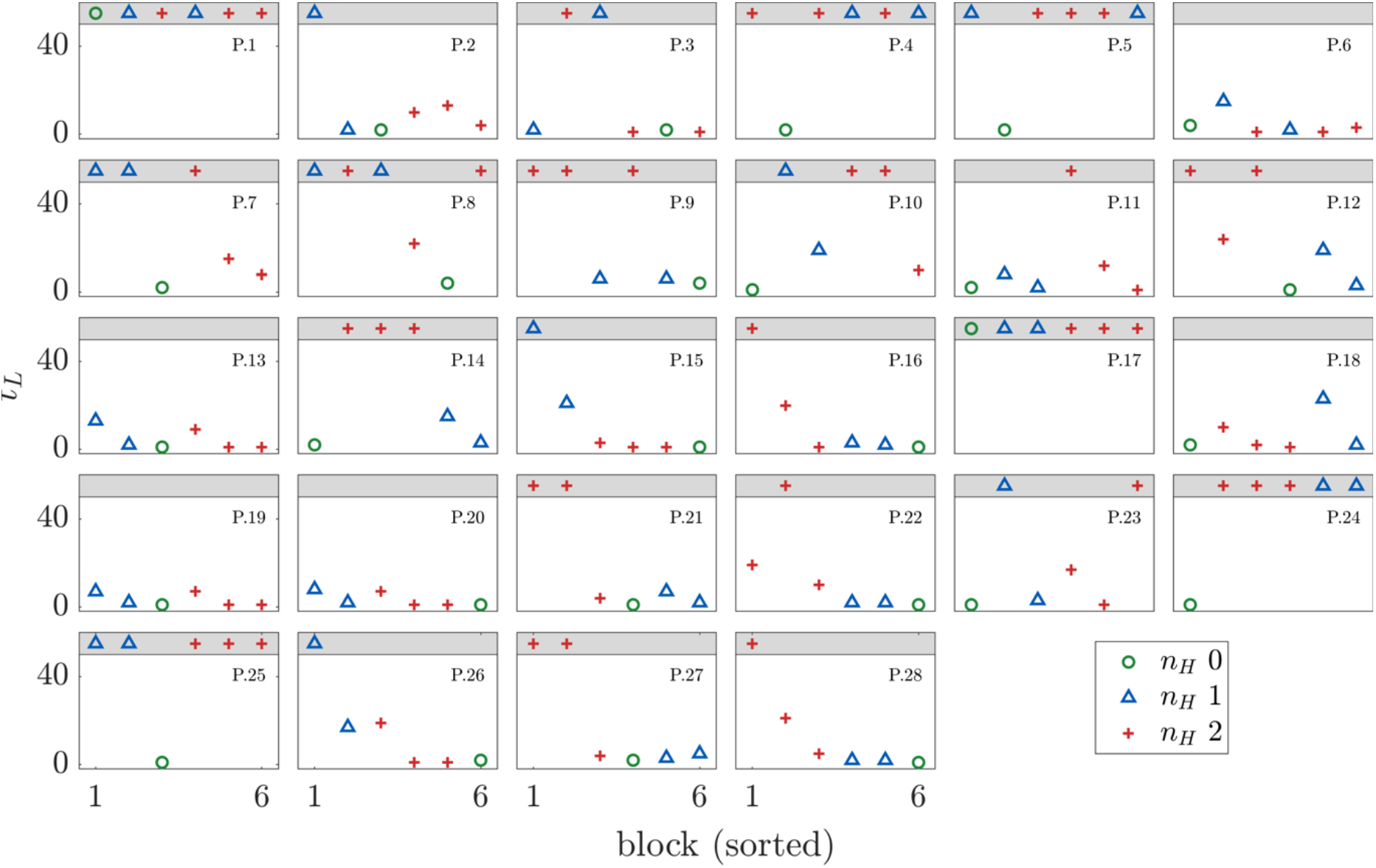
Learning time per block of recording in order of execution. Each plot is a different participant. On the y-axis, the learning time is reported in terms of episodes (above 50 means no-learning). Different horizons are depicted with different markers and colors (nH 0 green circle, nH 1 blue triangle, nH 2 red cross). Two participants did not learn any horizon, 4 participants learned nH 0, but not either nH 1 or nH 2, and the remaining 22 learned at least one of nH 1, 2.

Interestingly, FigSupp 2 shows that most participants, when they had learned the optimal strategy in one horizon, they then generalized and learned the optimal strategy in the next horizon much quicker. This collective result is presented in FigSupp 3, here we excluded the 2 participants who did not learn *n_H_* 0. The learning time is shown on the y-axis, each green dot refers to a participant, and the three plots refer to different horizons. In other words, the following plots show the learning time for each of the three horizons separately. Moreover, they show the difference if that specific horizon was performed before or after other conditions. Namely, panel (a) depicts the learning time in *n_H_* 0 in the case it was the first block carried out, or not. If *n_H_* 0 was performed during the first block, participants took a little longer to learn the optimal strategy than if they had already performed *n_H_* 1 or 2 before. Panel (b) illustrates the learning time in *n_H_* 1, separating the participants according to if *n_H_* 1 was performed before or after *n_H_* 2. Finally, panel (c) portrays the learning time in *n_H_* 2, separating the participants according to if *n_H_* 2 was performed before or after *n_H_* 1. Note that these plots show all participants, including the ones that never learned *n_H_* 1, 2, but did learn *n_H_* 0. From the results in (b-c), we can conclude that, in the horizons with consequence (i.e., *n_H_* 1, 2), participants needed less time to learn the optimal strategy of one horizon, if they had already performed the other one. We speculate that once the optimal strategy in any consequential block was understood, participants generalized the rule and by abstraction applied it to the other horizon.

**FigSupp 3.**
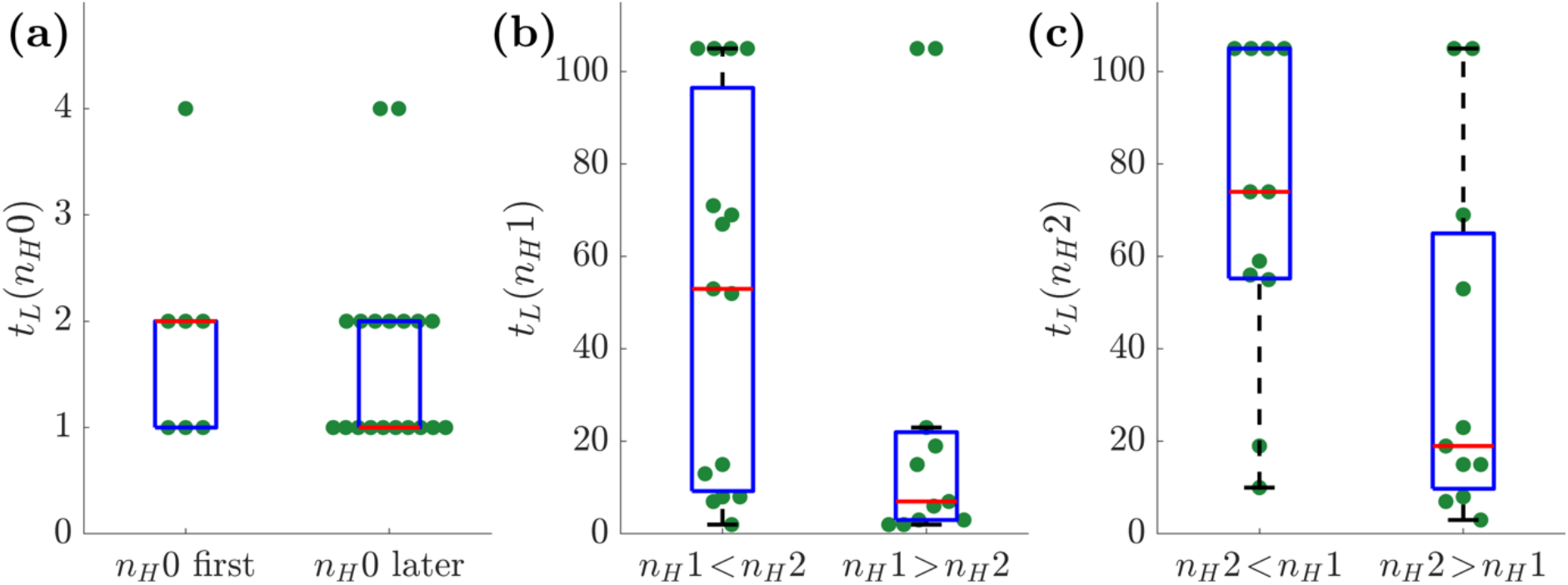
Learning time as a function of the horizon, and the order of execution. The 2 participants who did not learn nH 0 were excluded from this analysis. Green dots refer to participants, and the three panels refer to different horizons. **(a)** learning time in nH 0 in the case it was the first block carried out (left), or not (right). **(b)** Learning time in nH 1, for the participants that performed nH 1 before (left) or after (right) nH 2. **(c)** Learning time in nH 2, for the participants that performed nH 2 before (left) or after (right) nH 1.

### Parameter space analysis

To help reading Figure 7, FigSupp 4 shows the dependency of RT with the free parameters of Eq. 3. When not varied for the plot, we fixed the parameters to 1=67, =22 Hz, and α=0.001 in panels (a-c), and τ=39, =26 Hz, and α=0.005 in panels (d-f). Both the mean and standard deviation increase consistently with both the time constant τ and the threshold. The noise intensity σ does not have a substantial influence on RT.

**FigSupp 4.**
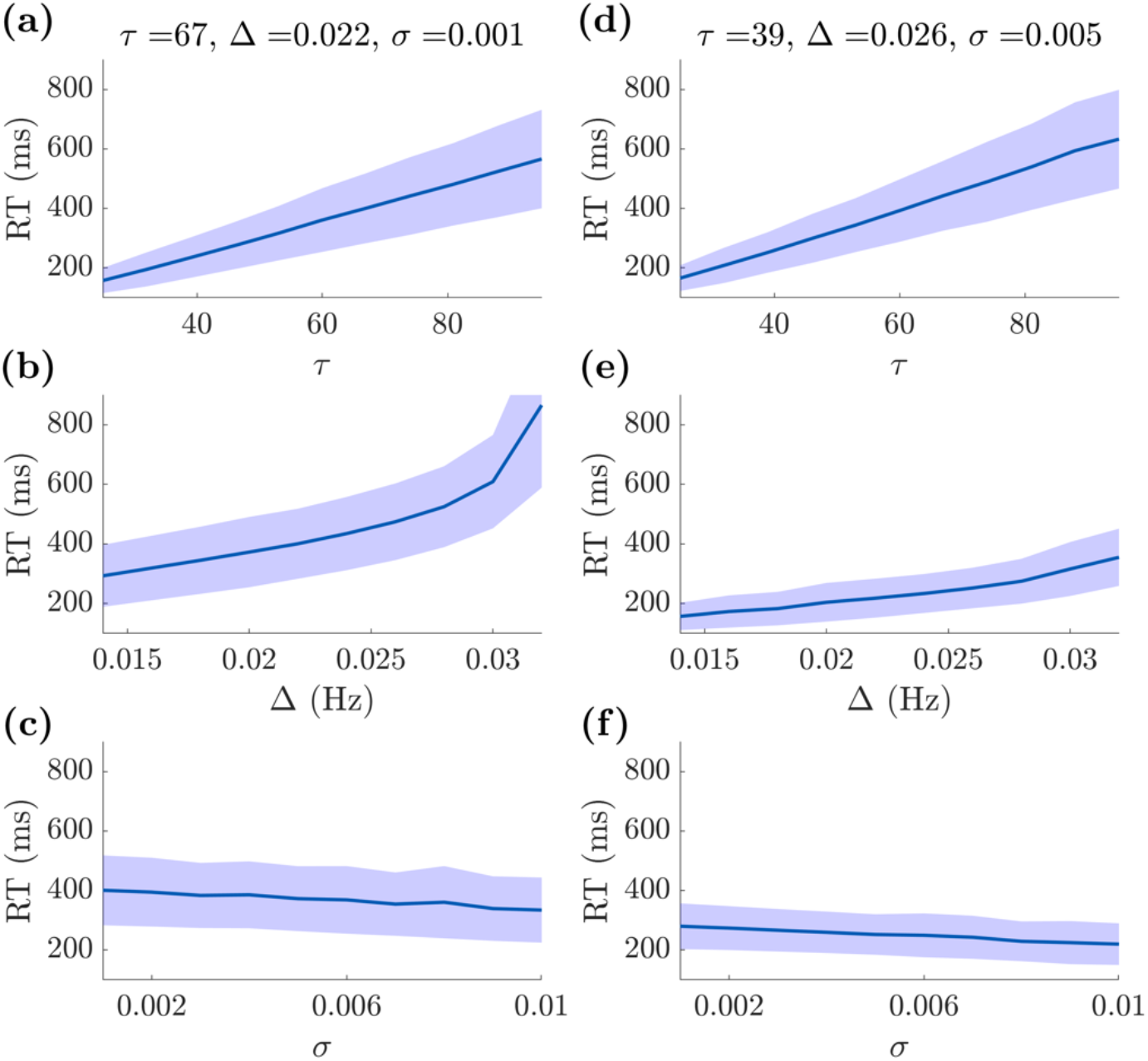
Parameter space analysis. Both the mean and standard deviation of the reaction time increase consistently with both **(a,d)** the time constant τ and **(b,e)** the threshold /-. **(c,f)** The noise intensity σ does not have a substantial influence on the reaction time. – Parameters used: **(a-c)** τ=67, /-=22 Hz, and α=0.001; **(d-f)** τ=39, /-=26 Hz, and α=0.005, when not varied for the plot.

## Notes

### Competing Interest Statement

The authors have declared no competing interest.

### Summary of Updates

Significant changes to title, introduction and discussion. Added supplementary materials.

